# Selective Targeting of Kinesin on Lipid Droplets in the Liver Reduces Serum Lipids

**DOI:** 10.1101/2025.11.17.688766

**Authors:** Subham Tripathy, Archisman Mahapatra, Ojal Saharan, Hindol Chatterjee, Neelanjana Sengupta, Siddhesh Kamat, Sreelaja Nair, Roop Mallik

**Affiliations:** Department of Biosciences and Bioengineering, Indian Institute of Technology Bombay, Mumbai 400076, India; Department of Biology, Indian Institute of Science Education and Research, Pune 411008, India; Department of Biological Sciences, Indian Institute of Science Education and Research Kolkata, Mohanpur 741246, India

## Abstract

The liver controls plasma lipids by secreting lipid-rich very low density lipoproteins (VLDL) into blood. Inside hepatocytes in the liver, Lipid Droplets (LDs) are transported to the smooth Endoplasmic Reticulum (sER) by kinesin-1 motors, and then catabolized in the sER to supply lipids for VLDL assembly. LDs are the only cellular organelle bounded by a phospholipid monolayer, and are thus distinct from all other (bilayer-bounded) organelles. It is therefore plausible that a given protein can bind to the LD membrane using mechanisms that are completely different from all other organelles. Indeed, here we find that kinesin-1 uses its tail domain to bind LDs, but alternative mechanisms to bind other organelles. A peptide corresponding to kinesin’s tail domain therefore competes with, and removes kinesin-1 ***<u>selectively</u>*** from LDs with minimal effect on other organelles. Delivery of lipids for VLDL assembly is consequently reduced, causing a remarkable reduction of ∼50% of secreted lipids (triglycerides and cholesterol) in cell culture. We further develop Orally fed Egg-liposomes as a method to deliver kinesin tail domain peptide to the liver of Zebrafish. The peptide reverses diet-induced hyperlipidaemia in Zebrafish larvae and brings the larvae back to a normolipidaemic state, thus confirming the effectiveness of our method in a physiologically relevant *in-vivo* situation. Strikingly, the peptide causes no unwanted accumulation of lipids in the liver, no toxicity and no developmental or behavioural defects in Zebrafish. Using a peptide to displace proteins (e.g. kinesin) selectively from LDs provides a conceptually novel and radically different approach against hyperlipidaemia. This *monolayer-versus-bilayer* strategy can be potentially extended to target other LD-bound proteins that function as key regulators of Lipid metabolism.

## Introduction

Lipids are stored inside cells in the form of triacylglycerol (TG) and Cholesterol esters (CE) that make up the neutral-lipid core of lipid droplets (LDs). LDs also harbour lipid-synthesizing enzymes, lipases and vesicular transport motors revealing LDs as dynamic hubs where lipids are stored or consumed to channel energy for metabolic demands (1–4). LDs share many common phospholipids and proteins with the ER (5, 6), however, LDs are different in that their bounding membrane is a phospholipid monolayer (Fig 1A). The monolayer membrane has unique biophysical properties (2), causing specific proteins to be targeted to LDs in response to metabolic and immunological cues (7, 8). Unlike bilayer organelles, no specific protein-targeting machinery has been identified for LDs. Rather, proteins could bind LDs via packing defects that appear on the LD membrane when TG is transiently exposed to the cytosol (9). LD proteins may thus belong to two classes (3, 10). Class I proteins translocate from the ER to LDs, and bind LDs through a hairpin structure (e.g. GPAT4, DGAT2). Class II proteins translocate from cytosol to LDs and often form amphipathic helices upon interaction with LDs (e.g. perilipins, CCT1).

**FIGURE 1.**
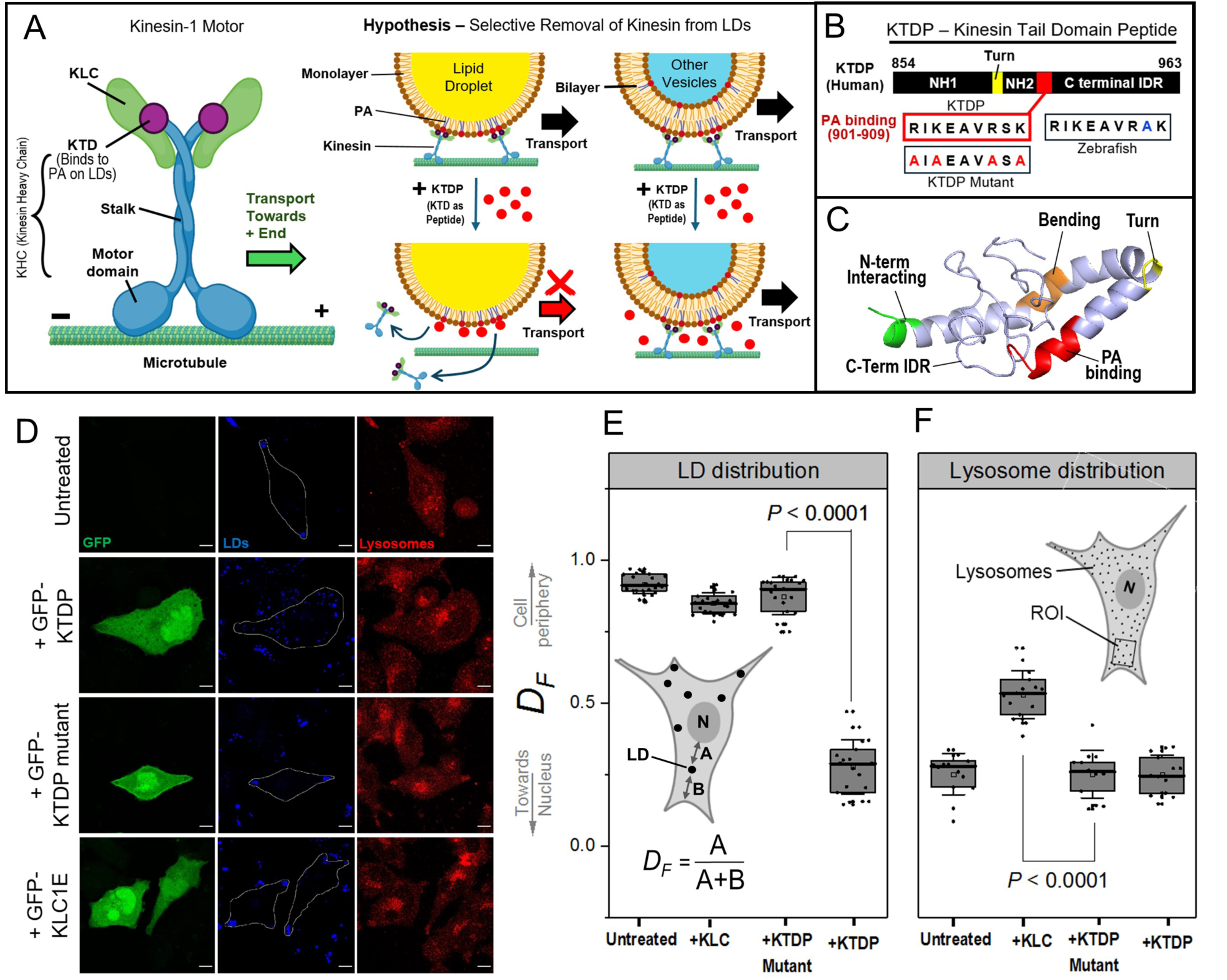
Selective Targeting of Lipid Droplet Transport Using a Kinesin Tail Domain Peptide (KTDP). A. Cartoon of the dimeric Kinesin-1 (Conventional Kinesin) Motor. Kinesin Heavy Chain (KHC) gene codes for the N-terminal Motor domain, Coiled coil stalk and C-terminal kinesin tail domain (KTD). Kinesin Light Chain (KLC) is expressed by the KLC gene. Two KHC and two KLC units combine to assemble the dimeric motor. **<u>Hypothesis</u>** depicts that kinesin-1 binds to lipid droplets (LDs) through its tail domain, thus a kinesin tail domain peptide (KTDP) can compete with and remove kinesin-1 from LDs. KTDP has minimal effect on other (bilayer) organelles which use alternative mechanisms (e.g. the KLC) to bind kinesin-1. Thus, LD transport can be targeted selectively to reduce delivery of lipids for lipoprotein assembly in hepatocytes. B. Details of human KTDP showing predicted N-terminal helices (NH1 and NH2), “turn” region, and a C-terminal intrinsically disordered region (C-Term IDR). The phosphatidic acid (PA) binding domain and its AA sequence are also shown. Mutations in basic AAs that block PA-binding of KTD (relevant to the KTDP-Mutant) are also shown. KTD sequence of PA-binding domain in Zebrafish is also shown. C. Predicted secondary structure of KTDP obtained from modelling in I-TASSER. D. Confocal images of McARH-7777 hepatoma cells (Untreated Control; Overexpressing GFP-KTDP, GFP-KTDP-mutant and GFP-KLC1E constructs). LDs (Blue : stained by MDH dye) and lysosomes (Red : stained using LAMP1 antibody) are shown in separate channels.Some cell outlines are marked in the LD channel. Scale bars = 10µm. E. Quantification of effects of overexpressing KTDP and KLC peptides on LDs using the parameter *D_F_* (see figure for definition). *D_F_* = 1 and *D_F_* = 0 respectively imply peripheral and perinuclear localization of LDs. Data represent Mean ± SD. F. Effects of overexpressing KTDP and KLC peptides on lysosome distribution. Lysosomes are quantified by measuring LAMP1 intensity within regions of interest (ROIs) near the cell periphery, and compared to LAMP1 intensity at whole-cell level. Ratio of peripheral to whole-cell LAMP1 intensity is then calculated (see main text). Each data point represents the distribution from an independent cell. Data are Mean ± SD.

Because these proteins hold the key for living organisms to access their energy/lipid reserves, the function of membrane-bound proteins on LDs is widely discussed in the context of LD biogenesis and metabolic disorders (10–12). Surprisingly, we find no discussion of the possibility that a protein’s presence on LDs could be manipulated against precisely these metabolic disorders. This is pertinent, considering the rising worldwide epidemic of Hyperlipidaemia and Obesity with very limited options available against these conditions (13–15). With this background, and because LDs are the only cytosolic organelle bounded by a monolayer membrane, we made the following **HYPOTHESIS** (See Fig 1A) :- Suppose that a given protein binds to monolayer or bilayer membranes using two different domains. A peptide that mimics the monolayer-binding domain of this protein should then compete with, and remove the protein from LDs with higher efficiency as compared to bilayer vesicles. This peptide could therefore inhibit LD-specific pathways downstream to that LD-resident protein with minimal interference in cellular functions of the same protein on bilayer membranes. To the best of our knowledge, such a possibility has never been discussed or demonstrated in the context of targeting LD biology and metabolic disorders.

Here we will test the above hypothesis for targeting the kinesin motor’s attachment to LDs versus other (non-LD) bilayer vesicles. Kinesin-1 is composed of two kinesin heavy chains (KHCs) and two light chain (KLCs) that are encoded by different genes and assembled into a dimeric motor (Fig 1A). The KHC consists of a motor, a coiled-coil stalk, and a C-terminal kinesin tail domain (16, 17). Kinesin-1 is recruited to many kinds of bilayer vesicles via protein complexes (e.g. Miro-Milton) and/or adaptors that interact with the KLC (18, 19). Kinesin-1 also binds to and transports LDs in *Drosophila* embryos (20) and in the rat liver (21, 22). We showed that insulin signalling activates phospholipase-D inside the liver to generate phosphatidic acid (PA) on LDs inside hepatocytes. Kinesin-1 then binds directly to PA, causing LD transport to the smooth-ER located at the periphery of hepatocytes (7, 22). At the smooth-ER, these LDs supply ∼70% of the lipid content of VLDL particles (23–25). Kinesin-1 knockdown reduced TG secretion from cells and also lowered serum-TG in animals (Rats) because kinesin mediated delivery of TG-rich LDs for VLDL assembly was reduced (22). Indeed, the peak of ApoB-100 (a VLDL marker) was shifted to higher density in sucrose gradient fractions, showing that TG-deficient VLDL particles of higher density were circulating in the blood of Rats after kinesin-1 knockdown (22).

While kinesin-1 knockdown did reduce serum-TG in animals, this cannot be a viable strategy to control plasma lipids because kinesin-1 has many other essential functions inside cells (18, 19). However, we found later that overexpressing the C-terminal kinesin tail domain (KTD; aa 854-963 of KHC – see Figs 1B and 1C) as a peptide disrupted **<u>only</u>** the peripheral distribution of LDs inside cells without affecting localization of lysosomes, early endosomes and mitochondria (7). To explain these findings, here we hypothesize that kinesin-1 binds to LDs via the kinesin tail domain (Fig 1A). Thus, the KTD peptide (hereafter KTDP) can competitively displace endogenous kinesin-1 from LDs to disrupt LD transport and reduce the lipid content inside VLDL particles. If KTDP is less effective in removing kinesin from other (bilayer) cargoes of kinesin, then cellular functions outside the LD-VLDL axis should not be perturbed significantly by KTDP. We support this hypothesis using *in vitro* assays, MD simulations and cell culture experiments. We further demonstrate that liposomes prepared from eggs (26) are a novel and facile *in-vivo* peptide delivery system to the liver of Zebrafish larvae and adults. KTDP is delivered intact to the liver, causing a marked reduction in serum lipids (TG and cholesterol) in the Zebrafish model which is widely used for studying lipid metabolism (27, 28). KTDP causes no adverse lipid accumulation in cell culture or in larvae, which continue to develop and behave normally. This strategy to reduce serum lipids by inhibiting kinesin-mediated delivery of LDs for VLDL assembly in the liver could provide a radically different approach against the growing epidemic of hyperlipidaemia.

## RESULTS

### KTD Peptide Specifically Disrupts Peripheral Distribution of Lipid Droplets inside Cells

We first tested the effect of overexpressing GFP-tagged KTDP on LD distribution in McA-RH7777 rat hepatoma cells which are known to secrete VLDL (7, 22, 29). We will use KTDP corresponding to the human KTD sequence throughout this work. Fig S1-A shows sequence alignment of KTD across the species relevant here (Human, Rat, Zebrafish). The helix region of KTD (aa 854-913 ; also see Fig S1-B) implicated in membrane binding (7, 30) is highly conserved, justifying the use of human KTDP in all our experiments. McA-RH7777 cells have an elongated morphology, with the smooth-ER located at the cell periphery where the plus-ends of microtubules (MTs) are present (7). Control (untreated) cells had most LDs at the cell periphery (Fig 1D), suggesting high kinesin activity on LDs. Overexpressing GFP-tagged KTDP (GFP-KTDP) caused LDs to redistribute throughout the cells, but a mutant GFP-KTDP (GFP-KTDP-mut; see Fig 1B) that cannot bind to PA on LDs (7) showed no effect on LD distribution (Fig 1D). We verified that GFP-KTDP and GFP-KTDP-mut are expressed at equal levels in cells (Fig S1-C), as also found earlier for these plasmids (7). We will also use GST and GFP-tagged versions of KTDP expressed in bacteria for our experiments (Fig S1-D). As described earlier (7), we quantified LD position using a fractional distance parameter (*D_F_* ; see Fig 1E). *D_F_* = 1 and *D_F_* = 0 indicate LDs respectively at cell periphery and cell-centre. GFP-KTDP cells had lower values of *D_F_* for LDs, but GFP-KTDP-mut caused no reduction in *D_F_* (Fig 1E). Importantly, GFP-KTDP had no visual or quantifiable effect on lysosome distribution in McA-RH7777 cells (Figs 1D, 1F). We have shown that KTDP also has no effect on the distribution of mitochondria and early endosomes in these cells (7). In contrast to KTDP, overexpressing GFP tagged kinesin light chain (GFP-KLC1E) had no effect on the distribution of LDs (Figs 1D, 1E). Contrastingly, GFP-KLC1E induced a peripheral distribution of lysosomes as also reported earlier (19).

Based on these observations, KTDP has a very pronounced and selective effect on kinesin-driven transport of LDs towards the peripheral region of cells. This selectivity can be explained if kinesin-1 binds to LDs using its KTD, but uses other domains to bind bilayer organelles (e.g. KLC for lysosomes). *In-vitro* studies show that the KTD binds to kinesin’s motor domain and also to the microtubule. KTD inhibits the ATPase activity to switch kinesin into an “off state” (31, 32). It is therefore intriguing that McA-RH7777 cells stably over-expressing KTDP show no phenotype and grow normally, with no effect on the localization of lysosomes, endosomes and mitochondria inside these cells (7). Why does the overexpressed KTDP not bind to the motor domain of kinesin-1 on bilayer organelles, and cause a general shutdown of kinesin-1 activity? This can be explained if association with KLC keeps kinesin-1 in an activated state that is immune to KTDP (32). We will return to these issues in the Discussion section.

### *In-vitro* Affinity of KTD Peptide to Monolayer and Bilayer Membranes

The above data suggest that KTDP inhibits kinesin-1 on LDs, but is unable (or less able) to inhibit kinesin function on other vesicles. The simplest explanation for this observation is that overexpressed KTDP competes with the KTD of endogenous kinesin-1 for binding to LDs, and thus removes kinesin-1 from LDs. KTDP neither removes nor does it block kinesin-1 function on other (bilayer) vesicles because ***(i)*** KTDP has lower affinity to bilayer membranes ***(ii)*** Kinesin-1 does not bind to bilayer membranes using KTD (Hypothesis; Fig 1A) and ***(iii)*** KTDP does not inhibit ATPase activity of kinesin-1 that is bound to bilayer vesicles because KLC keeps kinesin-1 in an activated state that is immune to KTDP.

We first tested for the removal of kinesin-1 by KTDP in a minimal reconstituted system using liposomes (these have a bilayer membrane) and artificial lipid droplets (ALDs) that have a monolayer phospholipid membrane around a TG core (7). Because KTD binds to phosphatidic acid (PA), liposomes and ALDs were both prepared using phosphatidylcholine (PC, 95%) and phosphatidic acid (PA, 5%). The ALD and liposome samples were normalized using the amount of PC detected on them by a thin layer chromatograph (Fig S1-E). For the same total surface area, a monolayer-LD should have approximately half of the PC content of a bilayer-liposome. Thus, we used appropriate dilutions to prepare a liposome sample with twice the amount of PC as compared to ALD sample, expecting these samples to have approximately same membrane surface available for KTDP binding (Fig S1-E). These samples were incubated with a GST-tagged KTDP prepared in bacteria (GST-KTDP; Fig S1-D), followed by centrifugation to separate ALDs and liposomes. Western blotting against GST demonstrated 4-fold higher intensity of GST-KTDP on ALDs as compared to liposomes (Fig. 2A). We will verify this preference of KTDP to monolayer membranes at single-vesicle level by optical microscopy in the next section.

**FIGURE 2.**
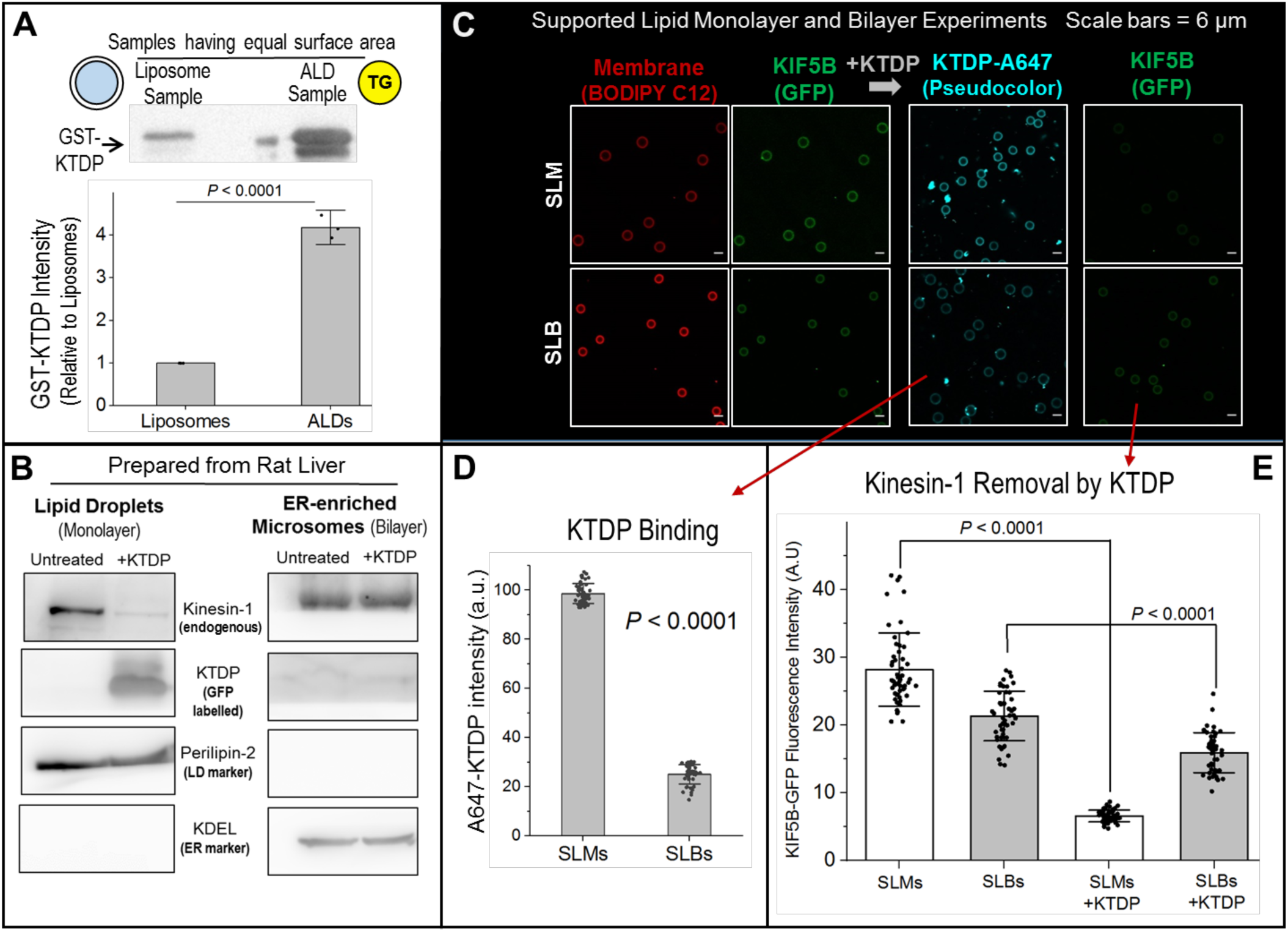
KTDP Preferentially Binds to and Removes Kinesin-1 from Monolayer Membranes *In-vitro*. A. Artificial LDs (ALDs) and Liposome samples are normalized to have approximately equal surface area (see main text). Samples are incubated with GST-KTDP, then separated from unbound GST-KTDP by cetrifugation. Immunoblotting with GST antibody shows ∼4-fold higher binding of GST-KTDP to ALDs. Data represent mean ± SD of three independent experiments. B. LD (monolayer) and ER-enriched microsome (bilayer) samples are prepared from rat liver. Each sample is divided into two equal parts. One part is left untreated and the other treated with GFP-KTDP. LDs or microssomes are separated from unbound GFP-KTDP by centrifugation. Immunoblotting against GFP shows significant binding of KTDP to LDs, but no detectable binding to microsomes. Perilipin-2 and KDEL immunoblots respectively show purity of LD and microsome samples. These immunoblots also show equal loading of LDs and microsomes across untreated and KTDP-treated lanes. C. Hydrophobic or hydrophilic beads (∼6 μm diameter) are incubated with BODIPY labelled liposomes (made from 95% PC + 5% PA). This results in formation of supported lipid monolayers (SLMs) or supported lipid bilayers (SLBs) with a fluorescent (BODIPY) membrane around them. SLMs and SLBs are then incubated with GFP tagged Kinesin-1 (GFP-KIF5B) to detect kinesin-1 recruitment on them in Confocal images. SLMs and SLBs are then treated with A647-labelled KTDP. KTDP-treated SLMs and SLBs are imaged to detect A647-KTDP and GFP-KIF5B on these structures. Scale bar = 6 μm. D. A647-KTDP fluorescence intensity on SLMs is significantly higher than SLBs, showing higher affinity of KTDP to SLMs. Each data point represents the integrated intensity of A647 fluorescence in a circular periphery around a single SLB or SLM. Error bars are Mean ± SD. E. GFP-KIF5B fluorescence intensity on SLMs and SLBs imaged before and after treatment with A647-labelled KTDP. Each data point represents the integrated intensity of GFP fluorescence in a circular periphery around a single SLB or SLM. Error bars are Mean ± SD.

### KTD Peptide Displaces Endogenous Kinesin-1 More Effectively From Monolayer Membranes

If KTDP binds preferentially to monolayers, then can it also displace kinesin-1 from LDs with higher efficiency as compared to bilayer organelles? To test this, we purified LDs and ER-enriched microsomes (i.e. bilayer vesicles) from rat liver. The LD sample and the microsome sample were each divided into two equal parts. One part was left untreated and the other was treated with GFP-tagged KTDP prepared in bacteria (Fig S1-D). LDs or microsomes were then separated from unbound proteins by centrifugation, and analysed by western blotting. GFP-KTDP was recruited abundantly to LDs, concomitant with significant loss of kinesin-1 from LDs (Fig 2B). Contrastingly, GFP-KTDP was barely detectable on microsomes, which retained almost all of their kinesin-1 even after KTDP treatment (Fig 2B). Equal loading of untreated and KTDP-treated LDs in western blot samples was verified by TG content (Fig S1-F). Fig 2B also shows western blots against perilipin-2 (LD marker) and KDEL (ER marker) which confirm the purity of LD and microsome samples. The similar intensities of perilipin-2 and KDEL shows equal loading of LDs and microsomes in untreated and KTDP treated western blot samples (Fig 2B).

The above described LDs/ALDs/microsomes are appropriate for bulk biochemical experiments, but their small size (<1 micron) precludes their use for optical imaging. To verify by direct imaging the effect of KTDP at a single-vesicle level, we developed an *in-vitro* assay as described in Fig S1-G. We used large (∼6 μm diameter) latex beads that were either hydrophobic (DVB-coated) or hydrophilic (Carboxylated). Liposomes with lipid composition (95%PC + 5%PA) were prepared and doped with a trace amount of fluorescent BODIPY-C12. These liposomes were then deposited onto the hydrophobic or hydrophilic beads to respectively create supported lipid monolayers (SLMs) or supported lipid bilayers (SLBs). BODIPY-C12 fluorescence imaging confirmed tight circular membranes on SLMs and SLBs (Fig. 2C). Fluorescence measured along a circular profile expectedly showed lower average intensity for the monolayer on SLMs than the bilayer on SLBs (Fig. S1-H). SLMs and SLBs were then adjusted by dilution to obtain samples that had the same optical density, and therefore the same number of beads/unit volume. We verified this normalization by visual counting of SLMs and SLBs using a hemocytometer. Because these two kinds of beads have approximately same diameter and they are present in equal number in the normalized samples, the total membrane surface area available for KTDP binding should be similar for SLM and SLB samples.

Cytosol was next prepared from cells overexpressing GFP-tagged kinesin-1 (GFP-KIF5B). SLMs and SLBs were incubated separately with cytosol to recruit GFP-KIF5B, then separated from cytosol via centrifugation. Confocal images showed that GFP-KIF5B was recruited to both SLMs and SLBs (Column 2, Fig 2C). These SLMs and SLBs were then treated with Alexa-647 labelled GST-KTDP, separated by centrifugation and imaged again. Alexa-647 signal on SLMs was significantly higher than SLBs (Column 3, Fig 2C). Quantification showed ∼4-fold higher Alexa-647 fluorescence per unit surface area on SLMs compared to SLBs (Fig 2D). This finding is in excellent agreement with the ∼4-fold higher KTDP detected on ALDs by western blotting (as compared to liposomes ; Fig 2A). Thus, KTDP has ∼4-fold higher affinity for the monolayer membrane, and might therefore be able to displace kinesin-1 more effectively from monolayers. To test this, GFP-KIF5B fluorescence was measured along the periphery of individual SLMs and SLBs (Fig 2E; at least 50 per condition). The below described ratios (*R_SLM_*, *R_SLB_*) were defined and measured from the images :-

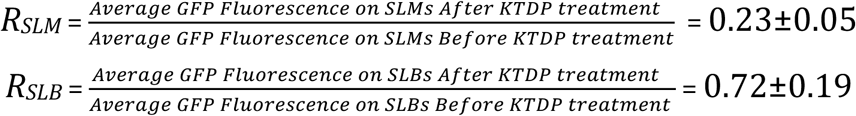

Thus, only ∼23% kinesin-1 was retained on SLMs after KTDP treatment (77% was removed). In contrast, ∼72% kinesin-1 was retained on SLBs (only 28% was removed). We used these ratios because the GFP signal on SLBs before KTDP treatment was slightly lower than SLMs (Figs 2C, 2E).

### KTD Peptide Reduces Lipid Secretion from McA-RH7777 Cells

If KTDP removes kinesin-1 efficiently from LDs, then does it also reduce lipid secretion from cells? To check this, we measured TG and CE secreted into medium from McA-RH7777 human hepatoma cells after overexpressing KTDP or a KTDP mutant that is defective for PA-binding (7, 30). These cells are well known to secrete VLDL (7, 22, 29). Both peptides were expressed at equal levels in these cells (Fig S1-C), as also seen earlier (7). Cells were first loaded with oleic acid (OA) to provide a substrate for LD formation. The OA-containing medium was removed, then cells were incubated in serum-free medium. The medium was removed by aspiration, and the cells were washed and harvested separately. LC-MS lipidomics of the collected medium showed that KTDP significantly reduced the secretion of specific long-chain TG species as compared to untreated and KTDP-mutant overexpressing cells (Fig. 3A and upper Inset, Fig 3A). Measurement using a commercial TG assay kit confirmed ∼50% reduction of TGs in the medium (lower Inset, Fig 3A). LC-MS also confirmed a drastic reduction of cholesterol esters in secreted medium (Fig 3B and upper Inset, Fig 3B). Measurement with a commercial cholesterol assay kit (detects both ester and free cholesterol) confirmed significant reduction of secreted cholesterol in medium upon KTDP overexpression (lower Inset, Fig 3B).

**FIGURE 3.**
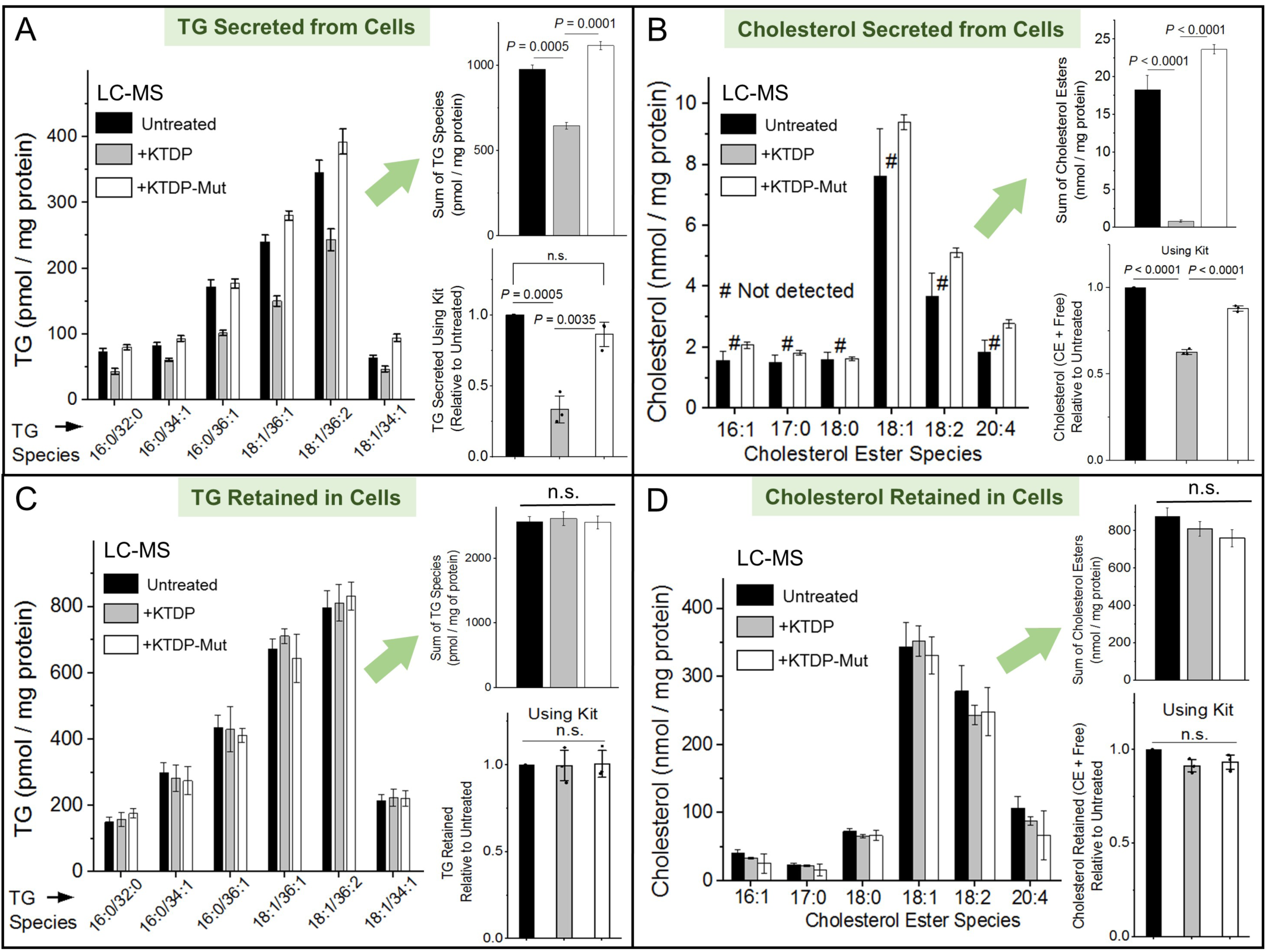
Effect of KTDP on Secretion and Retention of Lipids (McARH-7777 Hepatoma Cells). A. LC-MS for triglyceride (TG) in secreted media of untreated, GFP-KTDP and GFP-KTDP -mut overexpressing cells. The major species of detected TGs are shown. <u>Upper inset</u> shows the sum of detected TG species in LC-MS. Errors have been propagated. An overall reduction of ∼50% secreted TG is apparent from KTDP treated cells. <u>Lower inset</u> Similar reduction is also observed using a commercial TG assay kit. Error bars are SEM. B. LC-MS for Cholesterol in secreted media of untreated, GFP-KTDP and GFP-KTDP-mut overexpressing cells. The major species of detected cholesterol esters (expected to be present inside lipoprotein particles) are shown. Cholesterol ester secreted from KTDP treated cells was below the background levels in LC-MS data. <u>Upper inset</u> shows the sum of detected cholesterol ester species in LC-MS. Errors have been propagated. A drastic reduction of secreted cholesterol esters is apparent from KTDP treated cells. <u>Lower inset</u> Secreted cholesterol reduction is also observed using a commercial cholesterol assay kit that measures Esterified + Free cholesterol. Error bars are SEM. C. LC-MS for triglyceride (TG) retained inside untreated, GFP-KTDP and GFP-KTDP-mut overexpressing cells. The major species of detected TGs are shown. <u>Upper inset</u> shows the sum of detected TG species in LC-MS. Errors have been propagated. No significant difference in TG retention is apparent across conditions. <u>Lower inset</u> A TG assay kit also detects no significant difference across conditions. Error bars are SEM. D. LC-MS for Cholesterol retained inside untreated, GFP-KTDP and GFP-KTDP-mut overexpressing cells. The major species of detected cholesterol esters are shown. <u>Upper inset</u> shows the sum of detected cholesterol ester species in LC-MS. Errors have been propagated. No difference in cholesterol retained inside cells is observed across conditions. <u>Lower inset</u> A commercial cholesterol assay kit that measures Esterified + Free cholesterol also detects no difference across conditions. Error bars are SEM.

KTDP expression caused no increase in alanine aminotransferase (ALT) activity as compared to untreated or KTDP-mutant overexpressing cells, thus providing no evidence for KTDP-induced hepatocellular toxicity in McA-RH7777 cells (Fig S2-A). Very interestingly and surprisingly, LC-MS and commercial assay kit based measurements both showed that KTDP caused no increase inside cells of TG (Fig 3C and Insets of Fig 3C) or CE (Fig 3D and Insets of Fig 3D). Similar level of TG was detected inside untreated and KTDP-treated cells after a 12-hour pulse of Oleic acid, suggesting no significant effect of KTDP on LD biogenesis (Fig S2-B). How KTDP might reduce TG secretion without causing TG accumulation in cells remains open to future work. Preliminary experiments showed that the oxygen consumption rate (OCR) of KTDP treated McA-RH7777 cells was higher (Fig S2-C), which could arise from increased β-oxidation in mitochondria. LC-MS revealed that KTDP also increases free fatty acids (FFAs) retained inside cells (Fig S2-D), but causes no change in FFAs secreted from cells (Fig S2-E). It is therefore possible that KTDP induces a lipolytic pathway causing TG to be broken down into FFAs, which accumulate transiently inside cells before they are used up for β-oxidation. The exact mechanisms that prevent TG/CE accumulation inside KTDP-treated cells await further investigation. Nevertheless, if indeed this finding holds true *in-vivo* (will be tested in subsequent sections), then it has the exciting implication that circulating serum-TG and Cholesterol can be lowered by KTDP without the undesirable side-effect of fatty liver.

### KTD Peptide can be Delivered to Liver of Zebrafish Larvae Using Egg Liposomes

We next tested the effect of KTDP in a whole-organism model. We chose Zebrafish (*Danio rerio*), an established model for studying hyperlipidaemia and metabolic disorders (27, 28). The almost-transparent larvae permit *in-vivo* imaging of lipid distribution across visceral organs (e.g. liver, intestine) and the vasculature (26). Further, only a few species (Zebrafish, Humans, Rabbits, Hamsters) have an ortholog of the Cholesteryl Ester Transfer Protein (CETP) gene, a gene that is not found in rodents. Because CETP transfers neutral lipids between plasma lipoprotein particles (VLDL, HDL and LDL), the circulating lipoproteins in zebrafish closely resemble their human counterparts in composition and abundance (33). Zebrafish may therefore be more suitable than rodents to study the ill-effects of lipoprotein dysregulation. Indeed, atherosclerotic lesions are formed in Zebrafish kept on a high-cholesterol diet (34), but harder to observe in rodents.

Our objective was to test whether KTDP can reduce circulating lipids in zebrafish larvae. For the purpose of peptide delivery, we prepared liposomes using the yolk of chicken egg. Egg-liposomes have been used to deliver lipids and monitor lipid fluxes in zebrafish, they are also a simple method for simulating high-fat diet conditions in zebrafish (35). However, to best of our knowledge, egg-liposomes have never been used for peptide delivery to zebrafish. Peptides corresponding to human KTDP were used in Zebrafish as they are highly conserved (Fig S1-A). The PA-binding region of Zebrafish KTD is identical to humans except for a single S®A change (Fig 1B).

Three kinds of liposomes were prepared: No peptide (Empty), loaded with GFP-KTDP, or loaded with a GFP-KTDP mutant that cannot bind PA. The latter two conditions used 250 µg/ml concentration of peptide in the solution when preparing liposomes. Strong GFP signal appeared within nearly all liposomes of the last two categories, indicating successful peptide incorporation inside liposomes (Fig S3-A). These liposomes were labelled with trace amounts of BODIPY-C12, a fluorophore conjugated to a fatty acid chain that gets incorporated into TG and thus allows lipid visualization in larvae (33). Zebrafish embryos rely on yolk fats for nutrition until 5 days post-fertilization (5 dpf), after which they consume external food (27). We therefore used 6 dpf larvae for our studies. Liposomes were mixed into the water in which larvae were swimming (i.e. larval medium), with the expectation that larvae would eat up the nutrient-rich liposomes along with the KTDP inside liposomes. A schematic of the liposomal KTDP delivery experiments is shown in Fig S3-B.

Liposomes were administered in aforesaid manner over a 6-hour period with gentle agitation. Similar BODIPY-C12 signal was seen across different conditions immediately after this duration at the whole-larva level and inside the gut (Figs. S3-C). We therefore believe that larvae consumed the liposomes in similar quantity across conditions. We next asked whether the liposomes are actually helping in KTDP delivery to the larvae. GFP fluorescence was measured on serially diluted GFP-KTDP to prepare a calibration curve (Fig S3-D). We then measured GFP fluorescence in larval lysate prepared from larvae that had been fed with GFP-KTDP packaged inside liposomes, or GFP-KTDP simply mixed into water without any liposomes (again at 250 µg/ml concentration). Fig 4A shows the estimated KTDP concentration in whole larval lysate using the calibration curve of Fig S3-D. Barely any fluorescence was seen for KTDP without liposomes. Thus, the egg-liposomes do act as an effective peptide delivery agent to larvae. We next used transgenic zebrafish expressing the liver-specific marker FABP10a-mCherry to visualize the larval liver (Fig 4B). Liposomal delivery of GFP-KTDP or GFP-KTDP mutant caused robust GFP signal co-localising with FABP (Fig 4B), thus confirming KTDP delivery to the larval liver. As the GFP tagged KTDP is still fluorescent, the peptide appears to have been delivered to the liver without significant degradation.

**FIGURE 4.**
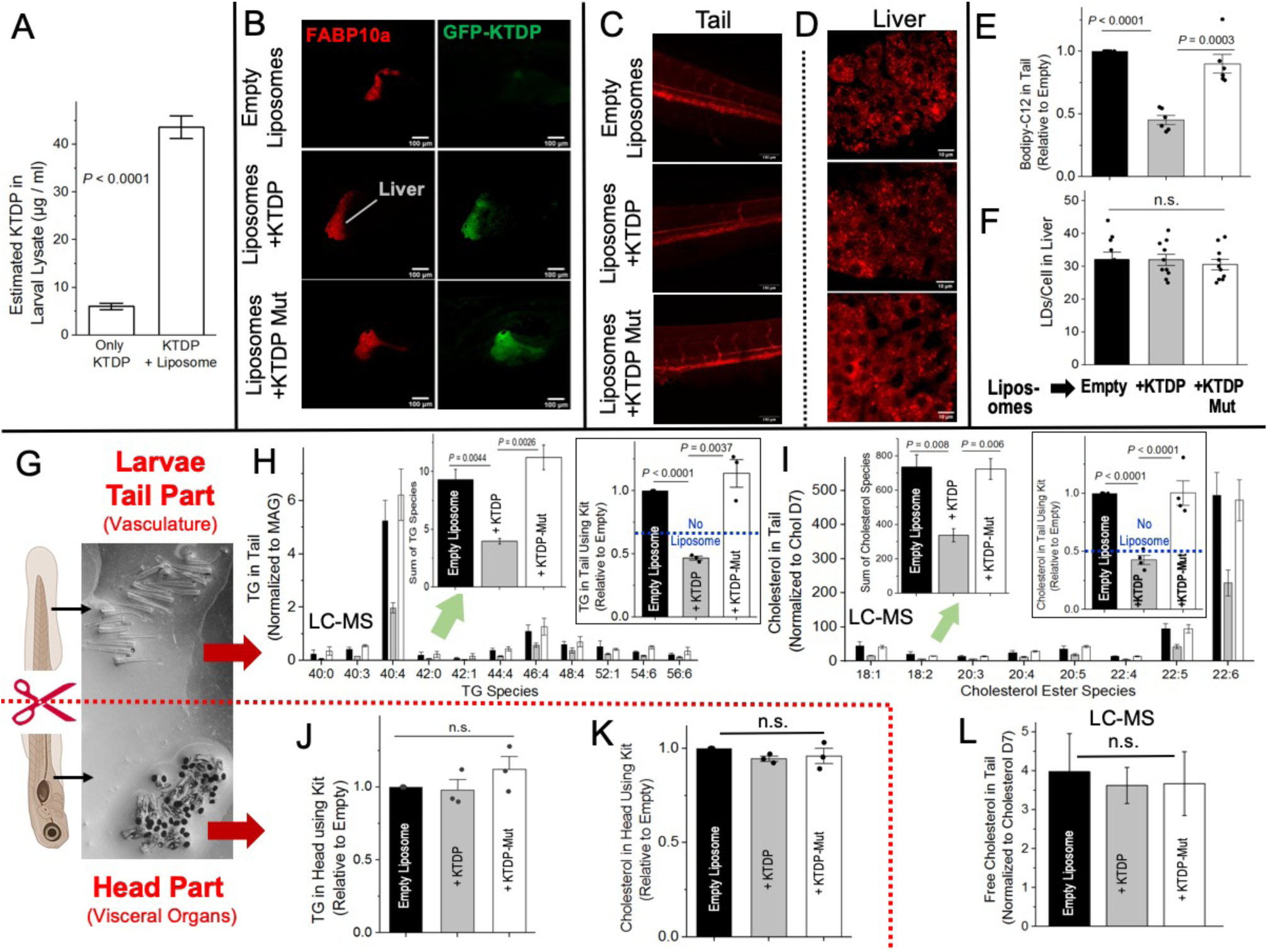
Liposomal Delivery of KTDP to Zebrafish Larvae : Effect on Lipid content in Tail and Head parts. A. Quantification of GFP fluorescence in zebrafish lysates prepared from larvae (6dpf) fed with GFP-KTD without liposomes (250 µg/ml), or GFP-KTD packaged in Egg liposomes. Calibration curve was used for estimating KTDP amount from GFP fluorescence (see main text). Error bars are Mean ± SEM. *N* = 3. B. Confocal images of zebrafish larvae after liposomal feeding of GFP-KTDP. Liver is marked in red in fish expressing FABP10a-mCherry (Red; liver marker). Scale bar = 100 μm. C. Confocal z-stacked images of the tail of larvae fed with BODIPY-C12 labelled liposomes of three types:- Empty liposomes, with KTDP or with KTDP-Mut. Scale bar = 100 μm. D. Confocal images of Nile Red staining in liver of larvae fed with liposomes as above. Scale bar = 10 μm. E. Quantification of BODIPY-C12 fluorescence in KTDP-treated larval tail (mean ± SEM, N=6). F. Counting of Nile Red stained LDs inside hepatocytes in larval liver shows no effect of KTDP (mean ± SEM, N=10). G. Larvae fed with liposomes (Empty, +KTDP or +KTDP-Mut) are bisected. The head (contains visceral organs e.g. liver, intestine etc.) and tail regions (vasculature) are separated and pooled for lipid estimation. We pooled 30 larval heads and 30 tails to make up a single replicate. Three such replicates were made for each condition of liposome treatment. Thus, a total of 30×3×3 (= 270) larval heads and 270 larval tails were used across the different replicates and treatments. H. LC-MS for secreted TG in pooled larval tails. **<u>Inset</u>**:- shows the sum of detected TG species. Errors have been propagated. An overall reduction of >50% secreted TG is apparent in KTDP-fed larvae. **<u>Boxed Inset</u>** :- Similar reduction is observed using a TG assay kit. The data are normalized to TG values for larvae fed with empty liposomes. Blue dashed line represents TG measured for untreated larvae (no liposomes; no KTDP). Error bars are SEM. I. LC-MS for secreted cholesterol in pooled larval tails. **<u>Inset</u>** :- Sum of all detected cholesterol ester species. Errors have been propagated. An overall reduction of >50% secreted cholesterol is apparent in KTDP-fed larvae. **<u>Boxed</u> <u>Inset</u>** :- Similar reduction is observed in the tail using a cholesterol assay kit. The data are normalized to cholesterol values for larvae fed with empty liposomes. Blue dashed line represents cholesterol measured for untreated larvae (no liposomes; no KTDP). Error bars are SEM. J. Measurement using a TG assay kit shows no effect of KTDP on TG content in head part of larvae (contains visceral organs). Error bars are SEM. K. Measurement using Cholesterol assay kit suggests that KTDP causes no change in Cholesterol in the head part of larvae. Error bars are SEM. L. LC-MS shows no effect of KTDP on Free cholesterol in tail part (vasculature) of larvae. Error bars are SEM.

### KTD Peptide Reduces Circulating Lipids in Zebrafish Larvae

Because the larval tail is devoid of visceral organs (28), BODIPY-C12 imaging in the tail allowed us to visualize lipids that had been secreted out into vasculature. KTDP reduced BODIPY-C12 fluorescence by ∼50% in the tail as compared to empty liposome or KTDP-mutant treated larvae (Figs 4C, 4E). Confocal imaging by Nile Red staining in the larval liver showed no accumulation or abnormal morphology of LDs or change in LD number after KTDP feeding (Figs 4D, 4F). We therefore believe that KTDP had no significant effect on LD biogenesis in the zebrafish liver. These findings are reminiscent of the unchanged TG and CE inside McA-RH7777 cells after KTDP expression (Figs 3C, 3D; Fig S2-B).

We next measured the TG in larval head part (contains visceral organs) and tail (contains vasculature) using biochemical methods. To do this, zebrafish larvae were fed with liposomes for 5 hours. Larvae were then washed, and individual larvae bisected to separate out the head and the tail parts (Fig 4G). We pooled 30 larval heads and 30 tails to make up a single replicate, and then measured TG and Cholesterol in each replicate. TG in KTDP-Liposome treated larval tail was reduced by ∼50% as compared to Empty or KTDP-Mutant liposomes, as seen by LC-MS (Fig. 4H) and also using a TG assay kit (Boxed inset, Fig 4H). Note that the lipid content of Empty-liposomes induces hyperlipidaemia as it increases serum TG by ∼40% compared to untreated larvae (no liposomes; blue dashed line in boxed inset of Fig 4H). Therefore, KTDP is able to restore normolipidemic conditions in lipid-fed hyperlipidaemic larvae. On similar lines, KTDP caused ∼50% reduction of cholesterol esters in larval tail (Fig 4I), but no change in free cholesterol in the tail (Fig 4L). These findings are in excellent agreement with the reduction of neutral lipids estimated by BODIPY fluorescence in larval tail (Figs 4C,4E). Again, reminiscent of the observation in McA-RH7777 cells, KTDP caused no increase of TG (Fig 4J) or cholesterol (Fig 4K) in the head part of the larvae.

### Time Course of Lipid Reduction, Phenotype, Mortality and Locomotion of Larvae after KTDP Feeding

How long does the lipid-lowering effect of KTDP last, and are there unwanted side-effects beyond this period? To explore, we fed 6 dpf larvae with liposomes that were Empty, filled with KTDP or with KTDP-Mutant. Larvae were fixed at 6,12,24,48 and 120 hours post liposome administration, then stained with Nile Red for imaging neutral lipids. We did not use BODIPY-C12 labelled liposomes because this dye gets metabolized and is not visible after 12 hours. Fig S4-A shows the staining and Fig S4-B quantifies change in Nile Red fluorescence at different time points. A significant reduction in lipids was seen in the tail (vasculature) of KTDP-fed larvae at 6 and 12 hours compared to empty or KTDP-mutant fed larvae, followed by return to baseline levels.

KTDP-fed larvae showed no observable deformity or abnormality upto 48 hours post-feeding as compared to empty or KTDP-Mut fed larvae (Fig S5-A). Mortality of larvae was similar across experimental groups upto 120 hours post KTDP feeding (Fig S5-B). To investigate potential long-term effects on larval locomotion, we stimulated larvae at the centre of a petridish with a micropipette tip 120 hours after KTDP feeding. We then measured where larvae first stopped after stimulation across concentric zones (Fig S5-C). No significant difference in first-stop zone was observed across conditions (Fig S5-D). Free-swimming larvae were also imaged 120 hours after KTDP feeding, and their motion tracked over six minutes (Fig S5-E). Analysis of tracks showed no differences in total distance travelled (Fig S5-F), or angular velocity during motion (Fig S5-G) across conditions. To sum up, we could observe no obvious effect of KTDP on the phenotype, mortality and locomotion of zebrafish larvae.

### Oral Delivery of KTDP to Adult Zebrafish: Effects on Serum and Liver Lipids

Because oral consumption is a preferred method for drug delivery, we attempted to deliver egg-liposomes containing KTDP to adult (1 year old) Zebrafish by oral gavaging. A gavaging procedure to deliver infectious agents to zebrafish (36) was adapted to deliver KTDP packaged inside egg liposomes (5µl of liposomes per fish; see Methods). Egg liposomes labelled with BODIPY-C12 (Empty controls, or loaded with KTDP) were administered by gavaging for three consecutive days at a KTDP concentration of 250 µg/mL, identical to the dose used in larval studies. Fish displaying bleeding, distress, or abnormal behavior were excluded from further experimentation. Fish were sacrificed four hours after the final administration. The tail (vasculature) was dissected out and centrifuged to collect blood and serum. The liver and anterior gut tissues were also dissected out for imaging and biochemical studies.

Fig 5A shows the gut and liver tissues for fish gavaged with empty liposomes. BODIPY signal in these tissues confirms that egg liposomes are delivered to the intestine and liver via oral route. Fig 5B shows the corresponding tissues for GFP-KTDP containing liposomes, where the GFP fluorescence suggests that KTDP is delivered intactly to these tissues. Figs 5C and 5D show a KTDP-induced reduction of TG and Cholesterol in serum of adult fish as compared to Empty Liposomes. Each data point in these figures corresponds to a single adult fish. Figs 5E and 5F confirm that KTDP causes no abnormal accumulation of TG or cholesterol in the liver of adult zebrafish, in agreement with similar observations in cell culture (Fig 3) and larvae (Fig 4).

**FIGURE 5.**
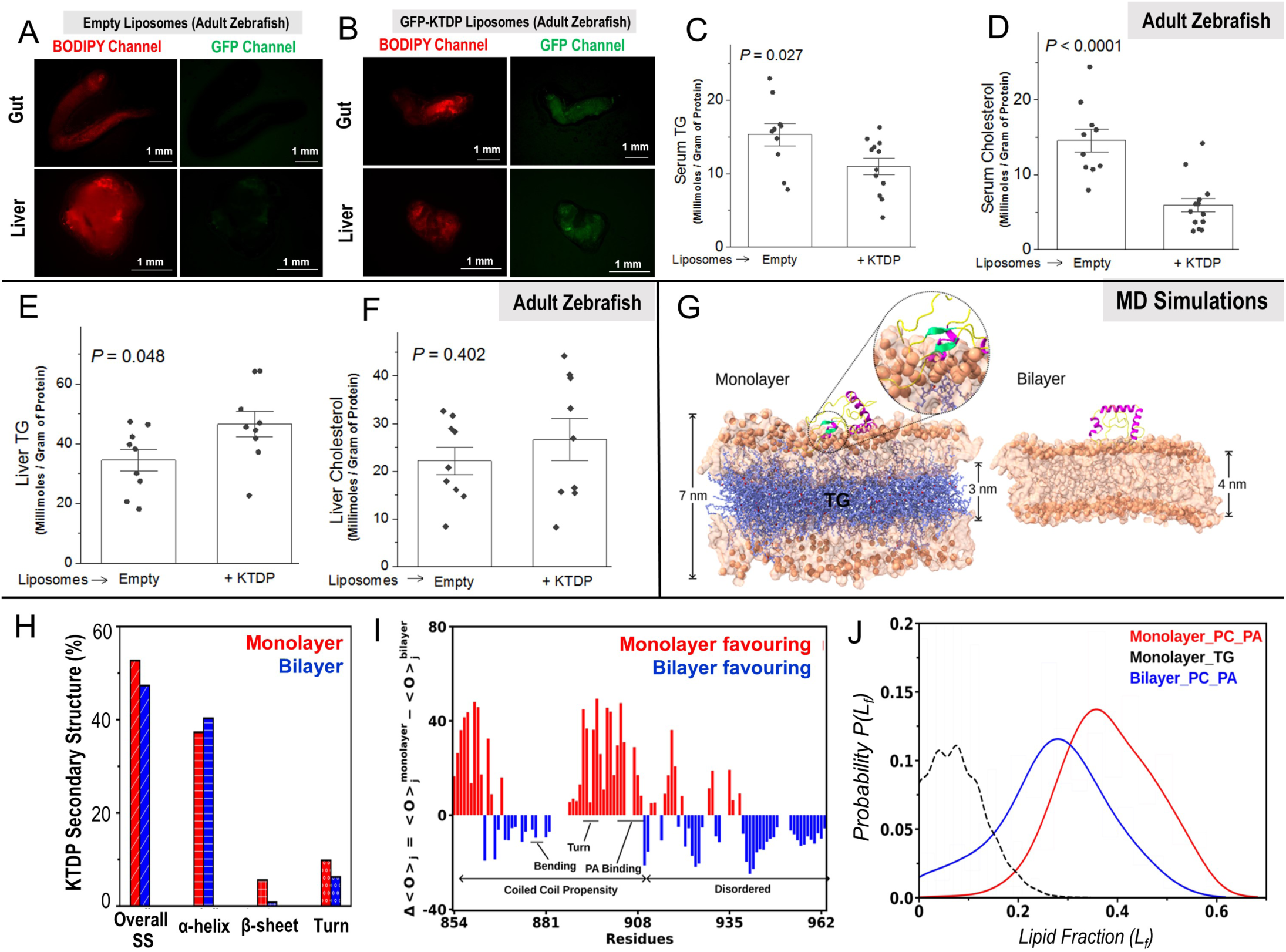
Lipid-lowering Effect of KTDP in Adult Zebrafish and MD Simulation of KTDP-Membrane Interactions. A. Confocal image of the Gut (intestine) and Liver dissected out from 1-year old Zebrafish. The fish were fed for 3 days with Empty egg-liposomes (containing no peptide) by an Oral gavaging method. Liposomes were labelled with trace amounts of BODIPY-C12 for visualization. B. Same as A, but with liposomes containing GFP-KTDP. GFP signal co-localizing with BODIPY suggests that the peptide was delivered intact to gut and liver tissues of fish. C. TG in blood (serum) obtained from the tail part (vasculature) of adult fish that were treated with Empty or KTDP-containing liposomes. Measurement was done using a commercial TG assay kit. Each data point represents a single fish. Error bars are SEM. D. Cholesterol TG in blood obtained from the tail part (vasculature) of adult fish that were treated with Empty or KTDP-containing liposomes. Measurement was done using a commercial TG assay kit. Each data point represents a single fish. Error bars are SEM. E. TG in dissected liver of adult fish treated with Empty or KTDP-containing liposomes. Measurement was done using a commercial TG assay kit. Each data point represents a single fish. Error bars are SEM. F. Cholesterol in dissected liver of adult fish treated with Empty or KTDP-containing liposomes. Measurement was done using a commercial cholesterol assay kit. Each data point represents a single fish. Error bars are SEM. G. Representations of KTDP-monolayer and KTDP-bilayer conformations in MD simulations. The upper and lower leaflets (translucent, orange), phosphorus atoms (orange beads), TG (purple), and KTDP secondary structures (magenta helix, yellow turns, green β-sheets) are shown. The aqueous phase is not shown for clarity. The average membrane thicknesses are reported in nanometers. H. Average secondary structure contents of KTDP in monolayer and bilayer systems. I. Difference in the lipid occupancy (Δ*O_j_*) of KTDP residues between monolayer and bilayer systems (see main text). The values on Y axis denote the time fraction of the entire simulation (expressed as a percentage) that a given residue spends in the vicinity of phospholipids (DOPC or DOPA). The calculation did not distinguish between headgroups versus tails of DOPC or DOPA. Specific regions of KTDP relevant to membrane binding are mentioned. J. Probability distribution of the Lipid fraction (*L_f_*) around monolayer and bilayer favouring residues of KTDP (see main text).

### Structural Changes in Kinesin Tail Domain Upon Membrane Binding

As shown above, KTDP can remove endogenous kinesin-1 from LDs to reduce TG/Cholesterol secretion from cells. We therefore explored the structural aspects of KTD-membrane interactions. Although crystal structures are not available, earlier analysis suggests significant coiled coil structures that terminate between AA 900-910 of KTD (37). Modelling in I-TASSER (Methods) predicted that KTD contains two N-terminal helices (NH1 and NH2) and a C-terminal intrinsically disordered region (C-IDR), as shown in Figs 1B and 1C. The helix region (aa 854-913) of KTD is highly conserved and therefore likely to be functionally relevant. A helical wheel representation of this region using Heliquest (38) showed accumulation of hydrophobic and hydrophilic amino acids on opposite sides (Fig S1-B). It is therefore possible that a conserved function of KTD is to recruit kinesin-1 to LDs by forming an amphipathic helix on the membrane (3, 10). Notably, the NH2 domain of KTD also contains a sequence (AA 901-909) that is rich in positively charged residues and implicated for binding to negatively charged PA on membranes (Fig 1B; Fig S1-B).

With this information, we undertook molecular dynamics (MD) simulations of KTDP binding to PA-containing monolayer and bilayer membranes (Methods). Following earlier work (39), we created a monolayer model containing a 3 nm thick slab of TG between two leaflets of phospholipids that were composed of 1,2-dioleoyl-sn-glycero-3-phosphocholine (DOPC) and 1,2-dioleoyl-sn-glycero-3-phosphate (DOPA; phosphatidic acid) taken in a 95:5 ratio (Fig 5G). The bilayer model was made with the same phospholipid composition (Fig 5G). A top view of membrane leaflets of monolayer and bilayer membranes is shown in Figs S6-A. As seen earlier (39), the monolayer showed surface-exposed TG molecules that likely cause packing defects, as also reflected in the higher area per lipid (APL) for monolayers (Fig S6-B). KTDP was placed near the upper leaflet and MD trajectories of 1.25 μs generated to interrogate the interaction of KTDP with monolayer and bilayer systems (Fig 5G). The dynamics of KTDP interacting with monolayer and bilayer systems is shown in Supplementary Videos 1 and 2.

On average, KTDP acquired higher secondary structure (SS) content in presence of the monolayer (Fig 5H). Structural flexibility in the monolayer was corroborated by enhanced distortion in the helical region (Fig S6-C). The loss in helical content on monolayer was offset by the emergence of β-sheets and turn (Fig 5H). Backbone root mean square deviation (RMSD) of KTDP was increased in the bilayer, indicating greater fluctuations and a transient nature of KTDP-bilayer interactions compared to monolayer (Fig S6-D). We next calculated the lipid occupancy (*O_ij_*), i.e. the time fraction for which the *j*_th_ amino acid residue of KTDP contacts a lipid indexed by *i* (*i* = DOPC or DOPA) . Fig 5I shows the difference in lipid occupancy between monolayer and bilayer systems as a function of *j :-*

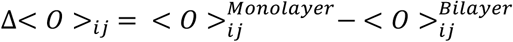

Here < *O* >*_ij_* is the averaged occupancy of *j*^th^ residue of KTDP across DOPC-DOPA (Methods). Only residues having occupancy >5% are reported. Positive (negative) values of Δ< *O_i_* > signify higher affinity of the *j*th residue to monolayer (bilayer) membranes. The average value of Δ< *O* >*_ij_* for monolayer favouring residues is ca. 23% and bilayer favouring residues is ca. 11%. Thus, overall KTDP-membrane interactions are higher for the monolayer system. We define the Lipid fraction as 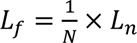 where *N* is number of KTDP residues interacting with *L_n_* lipids (DOPC or DOPA or TG) within a cutoff distance of 0.7 nm. The probability distribution *P*(*L_f_*) shows that residues favouring the monolayer interact more with lipids than residues favouring bilayer (Fig 5J). Considering only the non-zero lipid occupancy values, ca. 68% of the TG-interacting KTDP residues are also highly occupied with DOPC/DOPA (Fig S6-E), underlining the influence of TG in orchestrating lipid-KTDP interactions in the monolayer. In summary, the presence of interdigitated TG molecules enhances the APL of the monolayer membrane to create surface defects. The less compact membrane sustains interactions, causing SS gain and binding of KTDP to the monolayer.

## DISCUSSION

Elevated plasma lipids are associated with diabetes, cardiovascular disease and the metabolic syndrome. Reducing atherogenic plasma lipoproteins is the most accepted strategy against these ailments. Fibrates, omega-3 poly-unsaturated fatty acids (PUFAs) and Niacin are prescribed against hyperlipidaemia, but their ill-effects are well known (14, 15). Fibrates activate peroxisome proliferator-activated receptor-α (PPARα), a transcription factor regulating numerous genes in mitochondrial fatty acid oxidation. PUFAs down-regulate the SREBP-1c transcription factor and activate PPARα, they also stimulate LPL-mediated lipolysis and apoB degradation to increase VLDL clearance. Niacin inhibits adipose lipolysis, inhibits DGAT2 and reduces expression of apolipoprotein C-III for faster clearance of TG rich lipoproteins. Niacin however also causes liver damage and gastrointestinal problems. Statins effectively reduce cholesterol (and TG to some extent), but subjects still have a significant risk of cardiovascular disease requiring adjunctive therapy. Lomitapide, an inhibitor of the microsomal triglyceride transfer protein reduces TG assimilation into VLDL and chylomicrons, but also causes hepatic steatosis. Lipase inhibitors are known for many decades, but Orlistat is the only approved drug of this class. Orlistat provides only mild improvements in total cholesterol and lipoproteins (13). To sum up, most lipid-lowering therapies act via multiple known/unknown cellular pathways and have limited effect. Combination therapies are therefore often needed, thus amplifying their side-effects (14, 15). This situation underlines the need for novel management strategies against hyperlipidaemia.

The kinesin-dependent delivery of LDs for VLDL assembly has been discovered by us in recent years (7, 21, 22), and has therefore never been explored against hyperlipidaemia. The possibility that a peptide can remove an endogenous protein from the LD monolayer with minimal interference on bilayer vesicles is a conceptual advance that could be employed against other LD proteins. Particularly, other CYTOLD proteins that bind to LDs using amphipathic helices could be targeted. Kinesin driven LD transport for TG and cholesterol delivery towards VLDL assembly is demonstrated in human hepatoma cells, mouse primary hepatocytes and inside the rat liver (7, 22). Here we showed that KTDP can be delivered to the liver of zebrafish larvae and adults using egg liposomes, causing significant reductino of serum-TG and cholesterol (Fig 4, Fig 5). KTDP is part of endogenous kinesin-1, and therefore this peptide is unlikely to cause adverse immune responses even at higher dosage. Unlike the MTTP inhibitor Lomitapide that cause fatty liver, KTDP caused no increase of LDs in McA-RH7777 cells and zebrafish liver. While this is highly desirable, it remains to be understood why lipids are not accumulating in hepatocytes even when lipid secretion is reduced. KTDP had no effect on LD biogenesis because TG amount retained inside cells was unchanged (Fig 4C and Fig S2-B). Perhaps KTDP caused LDs to be targeted to mitochondrial oxidation and/or other degradative pathways, as evidenced by the increased oxygen consumption rate and FFAs inside KTDP treated cells (Figs S2-C,D). While these questions remain, it is reassuring to see no developmental, phenotypic or locomotory defects in zebrafish for significantly long periods after KTDP feeding (Fig S5).

Kinesin-1 not bound to cargoes inside cells may exist in a folded state where KTD binds to, and blocks the ATPase activity of kinesin’s motor domain (31). Accordingly, a KTD peptide also blocks ATPase activity of kinesin *in-vitro*. The over-expressed GFP-KTDP in our experiments is ∼1% of the total protein content of cells (see Fig S1-C). Even at such high concentration, why does KTDP have no effect on bilayer organelles? As demonstrated elegantly by others, KLC acts as an activator that prevents the KTD from inhibiting kinesin (32). KLC-mediated activation of kinesin on bilayer organelles may thus make them immune to KTDP. Because KLC overexpression had no effect on LD distribution (Fig 1E), KLC appears irrelevant to kinesin function on LDs. Further, PA is enhanced dramatically on LDs in the liver in fed state by the activity of phospholipase-D1 (7), causing vigorous kinesin-1 driven LD motion towards the smooth-ER (21). KTDP cannot bind LDs when charged residues in the PA-binding domain of KTDP are mutated (7). Thus, PA is essential for KTD/kinesin-1 binding to LDs. The generation of PA on hepatic LDs via phospholipase-D1 activity may be a signal for KTD binding and kinesin activation on LDs. Such a signal is likely absent on bilayer organelles, which use other mechanisms (e.g. KLC) to recruit and activate kinesin-1, and are therefore barely affected by the over-expression of KTDP.

Mutations within a 10-AA PA binding region (Fig 1B) render KTDP ineffective in blocking LD transport and lipid secretion (Fig 3 and Fig 4). Because this 10-AA region of KTD is critical for LD binding, shorter peptides around this region may also be effective in reducing lipid secretion. We modified the egg-liposome system for delivering the KTDP (with an additional GFP tag) to the liver of Zebrafish. The delivered KTDP was functional, as it reduced lipids very effectively in Zebrafish. KTDP was also possibly delivered to the intestine through ingestion of liposomes. Hence, the overall lipid reduction of ∼50% could arise via reduced TG/CE secretion from liver (in form of VLDL) as well as the intestine (in form of Chylomicrons). The effects of KTDP on TG/CE supply for chylomicron assembly, if any, remain to be explored. We note encouragingly that liver-targeted delivery of insulin using engineered liposomes is demonstrated (40). Engineered liposomes may further enhance KTDP delivery, and can be tested in rodents and primates. Small molecules mimicking KTDP may also serve this purpose.

To summarize, we earlier showed that kinesin-1 delivers LDs for assembling VLDL particles in hepatocytes. We hypothesized that kinesin-1 may bind LDs using KTD, but via alternate domains to bilayer vesicles. Thus, it is possible to remove kinesin-1 selectively from LDs by using KTDP as a competitive inhibitor (Fig 1A). Indeed, LD transport was inhibited when KTDP was delivered to human hepatoma cells, and zebrafish larvae treated with this peptide showed a remarkable reduction of ∼50% plasma TG and Cholesterol. Considering that the egg-liposomal delivery system works for delivering KTDP to liver of Zebrafish, we believe that KTDP and its peptidomimetic small molecules are an exciting avenue for exploration. Our results go beyond KTDP and its effect on lipid secretion. The possibility that endogenous proteins can be removed selectively from LDs to modulate metabolic (or other) processes has never been explored, and may be pursued for potential benefits in other pathways of lipid metabolism/LD biology.

## MATERIALS AND METHODS

### Reagents Used

GFP-KLC1E plasmid-a generous gift from Prof. mark Dodding, University of Bristol. DMEM (Sigma-D7777), DMEM no glucose (Gibco-11966025) FBS (Gibco-16000-044), trypsin-EDTA (15400054), Pen Strep (Gibco-15070063), Lipofectamine 2000 (Invitrogen-11668019), OPTI MEM (Gibco-31985070), Fatty acid free BSA (Sigma- A8806), BSA (Genei- 1650500501730), Monodansylpentane (MDH; abcepta-SM1000a), LipidTOX (Invitrogen- H34476), Silica gel 60 plates (Merck Milipore-1.05721.0001), Protease inhibitor cocktail (Roche-13539320), Ni-NTA sepharose resin (Cytiva-17531801), GST sepharose beads (Qiagen, 27-4574-01), egg PC (Avanti-840051P), egg PA (Sigma-P9511), Glycerol trioleate (Sigma- T7140), BODIPY C12 (Invitrogen-3822), Alexa 647 dye (Invitrogen-A20173), Chemiluminescent HRP substrate (Milipore-WBKLS0500), Latex beads, styrene divinylbenzene (Sigma-SD6A), Latex beads carboxylate (Polysciences-17141), Nile red (Sigma-72485), ALT assay kit (Cayman Chemical – 700260), TG assay kit (Elabscience - E-BC-K238), Cholesterol assay kit (Elabscience - E-BC-K109-S), Extracellular oxygen consumption assay kit (abcam-ab197243), Antibodies for Lamp1 (abcam-ab24170), KDEL (abcam-ab12223), perilipin-2 (Progen-651102), GFP (Invitrogen-A11122), α-tubulin antibody (Sigma-T9026), GST (Cloud clone-TAX158Ge22), secondary Alexa Fluor 555 donkey anti-rabbit (Thermo Fisher-A31572), donkey anti-rabbit IgG-HRP (Santa cruz-sc2313) and donkey anti-mouse IgG-HRP (Santa cruz-sc2314), Kinesin 1 (custom made).

### Statistics

Unless otherwise stated, error bars denote the SE of the mean (SEM). Unless otherwise stated, data were assumed to be normally distributed and Student’s *t*-test was used to calculate *P*-values for significance (Two-tailed ; 95% confidence ; Null hypothesis is that distributions are same). The *P* values have been mentioned at most places within figures. At some places following signs denote results of significance tests :- ***(*P* ≤ 0.001), **(*P* ≤ 0.01) and n.s. (*P* > 0.05).

### Animal strains and Procedures

All zebrafish (*Danio rerio*) care and experimental procedures were conducted in accordance with guidelines approved by the Institutional Animal Ethics Committee of IIT Bombay (Approval no. IITB/2025/BSBE/RM01). Zebrafish husbandry, breeding, and maintenance followed established standard protocols as described (41). Adult zebrafish, encompassing wild-type (Tübingen) and relevant transgenic lines [Tg(fabp10a:DsRed)], were maintained in a recirculating system with controlled water parameters [temperature (e.g., 28 ± 0.5°C), pH (e.g., 7.2-7.6), conductivity (e.g., 500-600 μS/cm)] under a 14-hour light / 10-hour dark photoperiod. Natural spawning was induced by placing male and female fish (typically in a 1:1 or 2:1 ratio) the evening prior into breeding tanks equipped with dividers. Dividers were removed the following morning upon commencement of the light cycle, and embryos were typically collected within 1-2 hours post-fertilization. Collected embryos were rinsed and subsequently reared in standard E3 embryo medium (5 mM NaCl, 0.17 mM KCl, 0.33 mM CaCl₂, 0.33 mM MgSO₄, buffered with 7.5 mM sodium bicarbonate to pH 7.3 and containing 0.0001% methylene blue) at ∼50 embryos in a 100 mm Petri dish in an incubator maintaining a constant temperature of 28.0 ± 0.5°C and the same 14:10 light:dark cycle. The E3 medium was refreshed daily to maintain water quality. Larvae were fed starting from 6 dpf, the feeding regimen consisted of a combination of dry feed and live artemia culture administered thrice daily. For procedures requiring immobilization such as imaging or fixing, larvae were anesthetized using 0.06% Tricaine.

Sprague-Dawley rats were bred and maintained by the animal house facility at the Tata Institute of Fundamental Research, Mumbai and animal protocols approved by the Institutional Animal Ethics Committee. Rats were maintained on a regular light (12-h)/dark (12-h) cycle and fed a standard laboratory diet. 8–12 week old male Sprague-Dawley rats were used for all the experiments.

### Cell lines and Growth Conditions

HEK-293T (ATCC CRL-11268) and McA-RH7777 (ATCC CRL-1601) cell lines were used. HEK-293T cells were cultured in DMEM supplemented with 10% fetal bovine serum (FBS). McA-RH7777 cells were grown in DMEM containing 20% FBS. All cells were maintained at 37 °C in a humidified incubator with 5% CO₂. Cells were subcultured using trypsin-EDTA upon reaching 70-80% confluency.

### LD and Lysosome Distribution in McA-RH 7777 Cells

McARH-7777 cells were seeded on acid-wash coverslips and allowed to grow till 50%-60% confluency. The cells were transfected with GFP-KLC, GFP-KTDP and GFP-KTDP Mutant plasmids using Lipofectamine 2000 (Thermo Fisher Scientific). Briefly, Plasmid DNA and Lipofectamine 2000 were separately diluted in Opti-MEM and Lipofectamine were mixed with Opti-MEM minimal media (Gibco) and then combined and incubated for 20 minutes at room temperature to generate transfection complexes. Control cells were treated only with lipofectamine, but no plasmid. The complexes were added to the cells and incubated for 6 hours followed by replacing the media with FBS (20%) containing high glucose DMEM media. Cells were grown for 48 h in the incubator and then treated with 400 µM BSA-conjugated OA for LD induction and cultured at standard conditions for 6 h. Cells were fixed with 4% PFA for 15 minutes and washed thrice with 1× PBS followed by permeabilization with 0.1% Triton X-100 in PBS for 10 minutes. The cells were again washed with PBS and blocked with 5% BSA in PBS for 30 minutes at RT. For lysosomal staining, cells were treated with primary LAMP1 antibody (1:100 dilution) for 1 hour at room temperature in a humidified chamber followed by washing thrice with PBS. The respective secondary antibodies (dilution 1:500) were added and incubated at RT for 1 h. The unbound antibodies were removed by washing with 1X PBS thrice. Cells were mounted with mounting medium and 50 µM monodansylpentane (MDH) for lipid droplet visualization. Fluorescence images were captured using a ZEISS LSM 900 confocal microscope with a 63× oil immersion objective. Image analysis was done using ImageJ-Fiji.

LD distribution was quantified by calculating the relative position of each LD within the cell (7). Lysosomal distribution was assessed by analyzing the spatial localization of LAMP1 fluorescence within the cell as discussed in (19). Fluorescence intensity in the peripheral region of the cell was measured and normalized to the total LAMP1 fluorescence intensity across the entire cell. The resulting ratio provides a measure of lysosomal distribution, with higher values indicating increased peripheral localization.

### Bioinformatics Analysis of KTDP

#### Secondary structure prediction, Multiple sequence alignment, Helical wheel diagram

Sequence alignment of KTDP across humans, rat and zebrafish was performed to assess evolutionary conservation. The analysis demonstrated a high degree of sequence identity with 94% similarity observed between human and rat KTDP, and 83% between human and zebrafish. The helical region critical for membrane binding (amino acids 854–913) exhibited strong conservation across all three species. To investigate the helicity nature of the N-terminal region of KTDP, a helical wheel projection was generated for amino acids 854-914 using HeliQuest web server (https://heliquest.ipmc.cnrs.fr/). The sequence was input in single-letter amino acid format, and an α-helical conformation was assumed.

#### Expression and Purification of GST-KTDP and GFP-KTDP from Bacteria

GST tagged fusion proteins were expressed in *E. coli* (BL21DE3) and purified using glutathione-conjugated sepharose beads using the manufacturer’s protocol. Briefly, BL21DE3 competent cells were transformed with pGEX-4T1 plasmid containing KTDP and KTDPmut fusion protein gene. A single colony was inoculated for primary culture at 37 °C overnight. Secondary culture (1% inoculum) was induced with 1 mM IPTG at OD600 = 0.6 and incubated overnight at 18°C with shaking. Cells were harvested by centrifuging the culture at 5000 g for 30 minutes and the pellet was resuspended in lysis buffer (50 mM TRIS-HCl, 150 mM NaCl, 5 mM beta-mercaptoethanol, 0.1% triton, 1 mM PMSF, 1 mg/mL lysozyme and 1X PIC, pH = 8.0) followed by lysis using probe sonication (amplitude=30%, 12 cycles, each of 15s ON and 45s OFF). Lysate was centrifuged at 20,000 rpm for 1 hour at 4°C and the supernatant was incubated with Glutathione Sepharose Beads pre-equilibrated with lysis buffer for 3 hours at 4 °C on a nutator. Beads were washed with 10 column volumes of wash buffer (50 mM TRIS-HCl, 150 mM NaCl, 5 mM β-mercaptoethanol, 1 mM PMSF, pH = 8.0) and bound proteins were eluted with reduced glutathione buffer (50 mM TRIS-HCl, 150 mM NaCl, 5 mM β-mercaptoethanol, 40 mM reduced glutathione, pH = 8.0). Eluate was dialyzed overnight against 1×PBS (pH = 7.4). Protein purity was assessed by 12% SDS-PAGE followed by Coomassie Brilliant Blue staining or anti-GST immunoblotting. Protein concentration was measured using Bradford assay.

GFP-KTDP-His proteins were expressed in *E. coli* BL21(DE3) using 1 mM IPTG induction at OD600 = 0.6 and incubation at 18°C, overnight. Cells were harvested by centrifugation followed by lysis using sonication in lysis buffer (50 mM sodium phosphate, 300 mM NaCl, 0.1% Triton X-100, 5 mM β-mercaptoethanol, 1× PIC, pH 8.0) and centrifuged (20,000 g, 1 h, 4°C). Supernatant was bound to Ni-NTA resin for 3 hours at 4^0^C on a nutator and washing was done with washing buffer (50 mM sodium phosphate, 300 mM NaCl, 50 mM imidazole). Protein were eluted with 300 mM imidazole in 50 mM sodium phosphate, 300 mM NaCl, 1X PIC. Eluted protein was dialyzed into 1X PBS and analyzed by SDS-PAGE/Coomassie.

#### Preparation of Artificial Lipid Droplets (ALDs) and Liposomes

ALDs were prepared by using a freeze-thaw technique (7). Briefly, 0.5 μmol egg phosphatidylcholine (PC) and 25 nmol of egg phosphatidic acid (PA) were mixed in clean glass tubes to prepare 1 ml of ALDs. This reaction mixture was dried under nitrogen gas stream for 30 minutes. Then 70 μl of glycerol trioleate was added in the dried glass tube followed by vacuum desiccation for 3-6 h to remove trace amounts of chloroform. After desiccation, 930 μl HKM buffer (50 mM HEPES-KOH, 120 mM potassium acetate, and 1 mM MgCl2, at pH 7.4) was added to the dried reaction mix for hydration. This reaction mixture was vigorously mixed by vortexing for 10 minutes to aid emulsification. The whitish emulsion was poured into ultra-low temperature resistant tubes and flash frozen in liquid nitrogen. The frozen emulsion was immediately thawed in a water bath preset at 55°C. This freeze–thaw process was repeated for five cycles with intermittent vortexing before every freezing step.

Liposomes were prepared by using a standard freeze-thaw technique (42). 1 μmol egg PC with 50 nmol of Egg-PA were dried in clean glass tubes and the reaction mixture was dried under nitrogen gas stream for 30 minutes followed by vacuum desiccation for 3-6 h to remove trace amounts of chloroform. The dried lipid film was hydrated with 1 ml HKM buffer (50 mM HEPES-KOH, 120 mM potassium acetate, and 1 mM MgCl2, at pH 7.4) and vortexed vigorously for 10 minutes. The mixture was subjected to five cycles of flash freezing in liquid N2 and thawing at 55°C. Each thawing step was followed by vigorous mixing to enhance unilamellar vesicle formation. For labelling the liposomes, BODIPY-C12 (8 μM) was added to the liposomes, mixed well and kept undisturbed at RT for 30 minutes.

#### Normalization of ALDs and Liposome Samples by Thin Layer Chromatography (TLC)

1 ml ALDs and 1ml liposomes were taken in two separate glass tubes. 2 ml methanol and 1 ml chloroform (2:1 v/v) were added to each tube followed by vigorous vortexing. The glass tubes were kept overnight at 4°C. The next day, 1 ml chloroform and water (1:1 v/v) were added to the tubes, vortexed, and kept undisturbed to allow phase separation under gravity (centrifuged at 1000 g for 10 minutes) to obtain a clear lower organic phase having phospholipids. Equal volumes of the lower organic phase were collected using Hamilton syringe and transferred to a new glass tube followed by drying under a stream of nitrogen gas (ReactiVap) for approximately 30 minutes. After drying, the phospholipid mix was resuspended in 50 ul of chloroform and spotted using a glass capillary on 8 X 10 cm Silica TLC plate (Merck) that was pre-rinsed with chloroform and dried overnight.

Phosphatidylcholine standards of amounts 12.5, 25, 50, 100 μg were loaded to generate a calibration curve. Phospholipids were resolved by using Chloroform: Methanol: Ammonium hydroxide (65:25:4) as solvent system and was run till the solvent front reached approximately 95% of the plate (43). The plate was air dried and the bands were visualized by spraying 10% CuSO4 in 8% H3PO4 followed by baking the plate in the oven at 100^0^C for 30 minutes. The plates were scanned and the bands were quantified using ImageJ. A calibration curve was generated using known phospholipid amounts to correlate band intensity with concentration. ALD and liposome phospholipid content was interpolated from the curve and normalized to surface area.

#### KTDP Interaction with ALDs and Liposomes

Normalized volumes (ALD:liposome = 2:1 v/v) of ALDs and liposomes were taken in two tubes. Equal amount (5 μM) of GST-KTDP protein were added to both the tubes and incubated on nutator at RT for 2 h. Both ALDs and liposomes were isolated from unbound proteins by density gradient ultracentrifugation using MEPS (5mM MgSO4, 5mM EGTA, 35mM PIPES pH=7.2, sucrose) buffer. Reaction mixture containing ALDs was supplemented with 1.5 times vol/vol of 2.5 M sucrose containing MEPS buffer and was loaded as the bottom layer of sucrose density gradient. This layer was overlaid with 5 ml (each) of 1.2M, 0.5M, and 0M sucrose in MEPS buffer. The gradient was centrifuged at 120000g using SW32 rotor at 4°C for 1 h to obtain lipid droplets (top-most whitish layer). Liposomes were pelleted at 100000g (transparent pellet). ALDs were collected using an 18-G needle and concentrated by centrifuging again at table top centrifuge at 20,000 g for 10 minutes. Liposome pellet was also resuspended in the same volume as concentrated LDs in 1X PBS.

#### Protein Precipitation and Western blotting

Equal volumes of ALDs or liposomes were mixed with chloroform:acetone (1:1 v/v) in two different safe lock tubes. The tubes are vortexed thoroughly and stored at -20°C overnight. The next day the tubes are vortexed vigorously and stored at -20°C for 15 minutes. Precipitated proteins were collected by centrifugation at 20,000 g for 2 h. The pellet was solubilized in 2× sample buffer and kept at 95°C for 15 minutes. The samples were loaded on 10% SDS-PAGE.

For Western blotting, the proteins on the SDS-PAGE gel were transferred to a PVDF membrane. The membrane was blocked with 5% non-fat dry milk in Tris-buffered saline with 0.1% Tween-20 (TBST) for 1 h at RT. The membrane was incubated with primary anti-GST antibody diluted in 5% BSA for overnight at 4°C. The membrane was washed thrice with TBST for 10 minutes each. The membrane was incubated with HRP-conjugated secondary antibody diluted in blocking buffer for 1 h at RT. The membrane was washed with TBST three times for 10 minutes each and developed using ECL. The blots were imaged on a ChemiDoc, and band intensity was quantified using ImageJ.

#### Isolation of Microsomes and Lipid Droplets from Rat Liver

Endoplasmic reticulum enriched microsomes were isolated using the protocol described in (42). Rats were anesthetized using sodium thiopentane (40 mg/kg body weight). Abdomen was cut open, and the liver was perfused with 50 mL of cold PBS (1× PBS) through the hepatic portal vein and then isolated. Liver tissue (9 g) was minced in cold room with 3 volumes of 0.25 M MEPS buffer containing 4 mM DTT, 8 μg/μL pepstatin, 4 mM PMSF, and Roche protease inhibitors. The minced tissue was then homogenized on ice in a 50-mL Potter-Elvehjem grinder using a ribbed Teflon pestle (up to 20 strokes). The homogenate was filtered through a two-layered cotton mesh and then centrifuged (8,700g, 4°C, 15 minutes) to obtain the postnuclear supernatant (PNS). The PNS was centrifuged (43,000g, 4°C, 7 minutes) to pellet mitochondria followed by centrifuging the supernatant (110,000g, 4°C, 60 minutes) to isolate the microsomes. Microsome pellet was resuspended in 1× PBS, flash-frozen, and stored at −80 °C for further experiments.

LDs were isolated from rat liver using the previously described protocol (7, 22, 42). Male Sprague–Dawley rats (3-4 months old) were anesthetized with sodium thiopentone (40 mg/kg, i.p.), liver was perfused with cold PBS via portal vein, excised, minced and homogenized in 1.5× (w/v) 0.9 M sucrose with MEPS buffer (with protease inhibitors) using a Dounce homogenizer at 4 °C. Homogenate was centrifuged (1800g, 10 minutes, 4 °C) to obtain the postnuclear supernatant (PNS) which was mixed with 1.5× vol of 2.5 M sucrose in MEPS buffer and layered under a step gradient of 1.2 M, 0.5 M, and 0 M sucrose in MEPS buffer. After centrifugation at 120,000g for 1 h at 4 °C, LDs (top whitish layer) were collected using an 18G needle, flash-frozen, and stored at - 80 °C.

#### Removal of Endogenous Kinesin-1 by KTDP from LDs and Microsomes

Isolated LDs were divided into two tubes having equal volumes. One tube was treated with 5 μM GFP-KTDP while the other tube received equal volume of 1X PBS as untreated condition. Samples were incubated on a thermomixer at 37^0^C for 1 hour at 500 rpm. Following incubation, unbound proteins were separated by sucrose density gradient ultracentrifugation (previously described). LDs were collected from the topmost layer and concentrated to equal volume. Treated and untreated LDs were normalized by optical density at OD₆₀₀ (22). Equal volumes of LDs were mixed with 0.1% SDS, heated at 60 degree for 30 minutes with intermittent vortexing, and centrifuged at 20000g for 20 minutes. The aqueous middle phase was collected, mixed with 1x loading dye and loaded on SDS-PAGE.

A similar protocol was followed for microsomes. Equal volumes of microsomes in separate tubes were incubated with 5 μM GFP-KTDP (KTDP treated condition) and 1X PBS (untreated condition) at 37^0^C for 1 hour with mixing at 500 rpm on a thermomixer. After incubation, unbound proteins were separated by ultracentrifugation and the microsomal pellets were resuspended with equal volumes of chilled 1X PBS. Equal volumes of treated and untreated microsomes were mixed with 1x loading dye and subjected to SDS-PAGE.

Western blotting was performed using anti-Kinesin-1 antibody to assess endogenous Kinesin-1 levels, anti-GFP antibody to detect GFP-KTDP binding, anti-Perilipin-2 as an LD marker, and anti-KDEL as an ER/microsome marker. Band intensities were quantified using ImageJ to compare endogenous Kinesin1 levels and GFP-KTDP binding between treated and untreated LDs and microsomes.

#### TLC for Normalization of LDs

Lipids from LDs were extracted by the methanol:chloroform method mentioned earlier. Extracted lipids were dried, resuspended in chloroform and then loaded onto silica plates pre-rinsed with chloroform. Separation was performed using two step solvent system i.e. in solvent I (n-hexane/ diethyl ether/acetic acid, 70:30:1) till half way and air dried, then in solvent II (n-hexene/diethyl ether, 59:1) for complete run. After air drying, TG bands were visualized by spraying the plate with 10% CuSO4 in 8% H3PO4, followed by baking >100 °C for 30 minutes.

#### Supported Lipid Bilayer (SLB) and Supported Lipid Monolayer (SLM) Preparation

6 μm carboxylated latex beads (hydrophilic) and 6 μm Divinylbenzene (DVB) coated latex beads (hydrophobic) were used for SLB and SLM preparation respectively. To avoid clumping, the beads were sonicated in a bath sonicator for at least 20 minutes. The beads were pelleted at 10,000 g for 5 minutes and then resuspended in milliQ water. For the preparation of SLBs and SLMs, 5 μl of carboxylated and DVB coated latex beads were added respectively with 1μl of 1M NaCl, 69 μl of autoclaved double-distilled water and 20 μl of the BODIPY-C12 labelled liposomes were taken together in a siliconized polypropylene microcentrifuge tube and kept for 30 minutes with intermittent vortexing (1 minute vortexing with 5 minutes rest). The mixture was washed thrice by the addition of 1 ml of milliQ water. SLBs and SLMs were pelleted at 10,000 rpm for 5 minutes and then finally resuspended in 200 μl of double-distilled water. The beads (carboxylated and DVB) were normalized to equal numbers using OD600 and manual counting using haemocytometer. BODIPY-C12 fluorescence was quantified on SLMs and SLBs by ImageJ to confirm the presence of monolayer and bilayer phospholipid membranes.

#### Preparation of KIF5B-GFP Enriched Cytosol from HEK-293 Cells

HEK-293T cells were transfected with pCDH-KIF5B-GFP plasmid. Cells were harvested 48 h post transfection, washed with ice-cold PBS, and resuspended in 3 ml of hypotonic buffer (10 mM Hepes [pH 7.4], 1 mM EDTA with protease inhibitor cocktail). The cells were incubated for 20 minutes in ice and centrifuged at 1200 g for 12 minutes. The cell pellet was resuspended in isotonic buffer (250 mM sucrose, 10 mM Hepes [pH 7.4], 1 mM EDTA, protease inhibitor cocktail), followed by lysis in a cell cracker (Isobiotec; 18-micron clearance). The lysate was centrifuged at 1200 g for 12 minutes to remove nuclei and cell debris. The resulting post nuclear supernatant (PNS) was centrifuged at 100,000 g for 1 h at 4^0^C and the supernatant (cytosolic fraction) is collected and stored at -80^0^C.

#### Labelling of GST-KTDP with Alexa 647

1 mg of the purified GST-KTDP protein was mixed with 50 ul of 1M sodium bicarbonate solution (pH = 8.5) and 0.5 ul of the Alexa Flour 647 (Invitrogen) dye in dark conditions. The mixture was incubated 2 hours at 4 °C on nutator followed by dialysis against 1X PBS overnight to remove the unbound dye molecules. Post dialysis, labelled A647-GST-KTDP proteins were collected and concentration was measured using Bradford assay.

#### KTDP Binding and GFP-Kinesin-1 Removal from SLMs and SLBs

Normalized volumes of SLMs and SLBs were incubated with equal concentration of cytosol containing KIF5B-GFP at 37^0^C for 1 h with intermittent mixing (5 sec on/5 sec off) in a thermomixer. KIF5B-GFP bound SLMs and SLBs were separated by centrifugation at 12000g for 10 minutess at room temperature and washed three times with 1X PBS to remove residual cytosol.

Collected SLMs and SLBs pellets were resuspended in 1X PBS followed by OD600 normalization. Equal volumes of normalized SLMs and SLBs were incubated with 5 μM A647 labelled GST-KTDP at 37^0^C for 1 hour with intermittent mixing. The second round of separation was done by centrifugation (12000 g, 10 minutes at RT). The pellets were resuspended in equal volumes of milliQ and imaged in confocal microscope. Fluorescence signals were quantified using ImageJ to compare A647-GST-KTDP binding and KIF5B-GFP removal between SLM and SLB.

#### Alanine Transaminase (ALT) Activity Test

To assess hepatocellular toxicity effect of KTDP, McARH-7777 cells transfected with GFP-KTDP and GFP-KTDPmut plasmids were measured for ALT enzyme activity using the ALT Colorimetric Activity Assay Kit (Cayman Chemical Cat# 700260).

Briefly, the cells were harvested and lysed to prepare cell lysate using RIPA lysis protocol. After centrifugation at 10,000 g for 15 minutes at 4°C, the supernatant was collected and kept on ice. For the assay 20μl of each sample was mixed with 150 μl of substrate and 20 μl of cofactor, and the plate was incubated at 37°C for 15 minutes. The reaction was initiated by adding 20 μl ALT initiator, and the absorbance was measured at 340 nm once in every 10 minutes up to 60 minutes. The assay and data analysis were done according to manufacturer’s instructions and compared across experimental conditions.

#### Oxygen Consumtion Rate (OCR)

Cellular respiration was measured using Abcam’s extracellular oxygen consumption assay kit (ab197243). Cells were seeded in a 96-well plate at a density of 20000 cells/well in 200 µL culture medium and incubated overnight at 37°C in a CO₂ incubator. Cells were transfected with the GFP-KTDP plasmid in four replicate wells, whereas control cells received no plasmid treatment. After 48 hours, oxygen consumption rate was measured following the manufacturer’s protocol. The culture medium was replaced with 150 µL fresh medium, followed by the addition of 10 µL reconstituted extracellular O₂ Consumption Reagent. Wells were sealed with 100 µL of pre-warmed high sensitivity mineral oil. Fluorescence was recorded at 2-minute intervals for 120 minutes at excitation/emission wavelengths of 380/650 nm. Fluorescence intensity/lifetime values were plotted against time, and the slope from the linear portion of the curve was calculated to determine the OCR.

#### Measurement of TG and Cholesterol Using Commercial Assay Kit and LC-MS

For measurement of secreted and cellular TG, both KTDP and KTDP-mut plasmids were transfected into McARH-7777 cells using the protocol explained earlier. 48 hours post transfection, cells were treated with 0.4 mM OA conjugated with BSA in incomplete media (no FBS) for 6 hours followed by washing with phenol-red-free incomplete media containing 0.5% fatty acid free BSA. For chase period, the cells were further cultured in phenol-red-free incomplete media with 0.5% fatty acid free BSA for 4 h. At the end of the chase period the media were collected and cells were harvested and stored in −80^0^C. Cell pellets were rinsed with cold 1X PBS and resuspended in RIPA lysis buffer supplemented with protease inhibitors. Cells were incubated on ice for 30 minutes with vortexing briefly in every 5-10 minutes followed by centrifugation at 14000g for 15 minutes at 4^0^C. The supernatant containing the cell lysate is collected. TG and total cholesterol measurement were conducted in secreted media and cell lysate using colorimetric assay kits (Elabscience) following the manufacturer’s protocol.

Secreted media and cell pellets were used for quantitative LC-MS analysis to measure TG, cholesterol and free fatty acids (FFA) following the protocol previously described in (22). Briefly, media and cell pellets (4 biological replicate for each group) were resuspended in 1 mL 1X-Phosphate buffered saline (PBS) and made up to a 4 mL mixture of 2:1:1 chloroform (CHCl3):methanol (MeOH): PBS. For semi-quantitative analysis of lipids, 1 nmol of an unnatural monoacylglycerol (C15:0 MAG) was added as internal standard for positive mode analytes. This homogenate mixture was vigorously vortexed and centrifuged at 3000g for 10 minutes to separate the mixture into an organic phase (bottom) and an aqueous phase (top) separated by a protein disk. The organic phase was removed by pipetting and stored on ice. To enhance the extraction of phospholipids from the aqueous layer, 100 μL of formic acid (MS grade, Honeywell, Catalog # 94318) was added, and this mixture was vigorously mixed. 2 mL of CHCl3 was added, and this mixture was vortexed and centrifuged as described previously. The organic layer was pooled with the one from the first extraction step and dried under a stream of nitrogen gas. The dried lipid extracts were re-solubilized in 200 μL of 2:1 CHCl3:MeOH and 10 μL was injected into an Agilent 6545 LC-QTOF (quadrupole-time-of-flight) LC-MS/MS for semiquantitative analysis using high-resolution auto MS-MS methods and chromatography techniques. LC separation employed a Gemini 5U C-18 column (Phenomenex) coupled with a Gemini guard column (Phenomenex, 4x3 mm, Phenomenex security cartridge). The buffers for positive ion mode runs consisted of 95:5 H2O: MeOH + 0.1 % Formic acid + 10 mM ammonium formate (buffer A) and 60:35:5 Isopropanol: MeOH: H2O + 0.1% Formic acid + 10 mM ammonium formate hydroxide (buffer B). Methods spanned 60 minutes, starting with 0.3 mL/min 100% buffer A for 5 minutes, 0.5 mL/min linear gradient to 100% buffer B over 40 minutes, 0.5 mL/min 100% buffer B for 10 minutes, and equilibration with 0.5 mL/min 100% buffer A for 5 minutes.

ESI-MS analysis settings included drying gas and sheath gas temperatures at 320 °C, a flow rate of 10 L/min for both drying gas and sheath gas respectively, a fragmenting voltage of 150V, capillary voltage of 4 kV, nebulizer (ion source gas) pressure set at 45 psi, and nozzle voltage of 1 kV. For analysis, a lipid library in the form of a Personal Compound Database Library (PCDL) was employed, and peak validation relied on relative retention times and fragments acquired. Quantification of all lipid species involved the normalization of areas under the curve to the corresponding internal standard area and further normalization to the total cell count. Subsequently, the changes were plotted for each lipid in comparison to the appropriate control within the group.

Cholesterol and cholesteryl esters were quantified using an Agilent 6545 QTOF mass spectrometer coupled with a 1290 Infinity II UHPLC system (44). Separation was performed on a Gemini C18 column using a 30-minute gradient LC method with formic acid and ammonium formate-containing solvents. MS acquisition was done in positive ion mode using Auto-MS/MS with optimized parameters and a preferred ion list targeting cholesterol and cholesteryl esters.

#### Preparation and Administration of Egg Liposomes to Zebrafish Larvae

Liposome preparation and feeding procedures were conducted following (35), with slight modifications. To prepare BODIPY-tagged liposomes, red fluorescently labelled BODIPY™ FL C12 (Invitrogen, Cat. No. D3822) was evaporated under a stream of nitrogen (N₂) and resuspended in 10 µL of 100% ethanol in Eppendorf tubes. Zebrafish embryo medium (EM; 90 µL) was then added to the resuspended solution. The resulting fluorescent stock solutions were protected from light and stored at 4°C.

For the liposome feeding solution, 1 mL of frozen chicken egg yolk (stored at -80°C) was thawed to room temperature and mixed with 19 mL of zebrafish EM. The peptides KTDP and KTDP-Mut were incorporated at this stage, ensuring a final concentration of 250 µg/mL in the feeding medium. The mixture was pulse sonicated for 40 s (1s on, 1s off; output intensity: 40%), filtered through a 40 µm cell strainer, and subjected to a second sonication step before collection. The resuspended BODIPY C12 was then added to 5 mL of the emulsion and vortexed for 30 seconds to achieve a final concentration of 6.4 µM BODIPY C12.

Freshly prepared liposomes were diluted tenfold and introduced into a flow chamber on a transparent glass slide. After allowing the liposomes to settle for 5 minutes to facilitate adherence to the slide’s bottom surface, imaging was performed using a confocal microscope (Zeiss LSM 900) equipped with a 63X objective. ESID and GFP excitation lasers were utilized to visualize liposome particles, confirming both the successful formation of liposomes and the encapsulation of KTDP.

Prior to liposome feeding, larvae were screened for any developmental abnormalities. A total of 15 healthy larvae per well were placed in 3 mL of the liposome solution and fed in a 6-well culture dish on an orbital shaker (30 rpm) for 6 hours across three experimental groups: BODIPY Liposomes (Empty), BODIPY-liposomes with KTD Wild type-GFP peptide (KTDP), and BODIPY-liposomes with KTD mutant-GFP peptide (KTDP-Mut). No observable signs of toxicity were detected during and after the feeding. Successful feeding was confirmed by the presence of red fluorescence in the gut area, observed under a fluorescence microscope (Olympus MVX10). To assess the delivery of the peptides specifically to the liver, a separate set of experiments was conducted using the same three experimental groups (without BODIPY-tagged liposomes) on transgenic zebrafish larvae [Tg(fabp10a:DsRed)], where liver cells express red fluorescence under the fabp10a promoter. After feeding in both sets of experiments, larvae underwent three sequential washes in fresh EM (5 minutes per wash) to remove residual liposomes or free peptides, followed by anesthesia using Tricaine. Larvae were then fixed in 4% paraformaldehyde for imaging or snap-frozen in liquid nitrogen and stored at -80°C for biochemical assays.

#### Efficiency of Liposome-mediated Peptide Delivery in Zebrafish Larvae

To evaluate the efficiency of liposomal delivery for GFP-tagged peptides (KTDP and KTDP-Mut), a standard curve was generated by plotting protein concentration against GFP fluorescence intensity. GFP excitation and emission wavelengths were set at 488 nm and 509 nm, respectively, with fluorescence measurements conducted using a multimode plate reader (Agilent Synergy H1) in a 96-well plate. Serial dilutions of KTDP-GFP were prepared at concentrations of 40, 50, 67, 100, 125, 200, and 250 µg/mL, and their corresponding fluorescence intensities were recorded. The resulting standard curve was used for quantification. Following peptide administration, 6 dpf zebrafish larvae were incubated with treatments for 6 hours in 6-well plates, as previously described. 50 Larvae were distributed per condition, covering three experimental groups: liposomes without GFP-KTDP, GFP-KTDP (250 µg/mL) encapsulated in liposomes, and GFP-KTDP (250 µg/mL) directly mixed into the media without liposomes.

Larvae were homogenized using probe sonication (2s on, 3s off; total 40s; 40% intensity), followed by centrifugation at 10,000 g for 10 minutes to collect the resulting supernatants. To account for background fluorescence, lysates from the group containing liposomes without GFP-KTDP were used as the negative control. The fluorescence intensity obtained from these samples was subtracted from all experimental groups to ensure accurate assessment of liposomal delivery efficiency.

#### Lipid extraction and Estimation of TG and Total Cholesterol in Larval Zebrafish

For the quantification of TG and total cholesterol, frozen zebrafish larvae were thawed on ice and bisected using a blade to separate the head region (containing visceral organs like liver, gut, brain etc.) from the tail region (primarily vasculature). The head and tail portions from 60 larvae were pooled to constitute one biological replicate, and this entire procedure was performed in triplicate (n=3). The pooled head tissues and the pooled tail tissues from the same 60 larvae (for one replicate) were each placed in 180 µL of PBS and sonicated using a pulse sonicator (2s on, 3s off; 50% intensity) for 30 seconds. The resulting homogenates were then centrifuged at 12,000 × g for 10 minutes at 4°C, and the supernatants were collected separately for head and tail regions.

Lipids were extracted from the supernatants of both head and tail homogenates using the chloroform-methanol method. Briefly, the collected supernatants were completely dried in a vacuum concentrator under constant rotation at room temperature. Once dried, lipids were extracted by adding a chloroform-methanol (2:1) mixture to the dried supernatant, followed by centrifugation at 12,000 × g for 15 minutes. The organic phase containing chloroform was collected and dried under a nitrogen (N₂) stream. The dried lipids were then reconstituted in 50 µL of isopropanol and stored at -20°C until further use. Protein concentration was determined using Bradford’s method from the initial aqueous supernatants of both head and tail homogenates for each replicate. Prior to TAG and total cholesterol measurements, the resuspended lipid extracts were normalized for protein content. The same tail lysates were subjected to lipid extraction and lipidomics analysis as described above.

TG and total cholesterol quantification was performed using colorimetric assay kits (Elabscience, Cat No. E-BC-K238 and E-BC-K109-S respectively) following the manufacturer’s protocol. Briefly, the protein-normalized lipid extracts were mixed with the respective kit’s reagent, incubated at 37°C for 30 minutes, and the absorbance was measured at 510 nm using a microplate reader (Agilent Synergy H1).

#### Imaging and Quantification of Circulatory Lipids in Zebrafish Larvae

Zebrafish embryos, following experimental treatments, were fixed in 4% paraformaldehyde (PFA) overnight at 4°C. To remove residual fixative, the embryos were washed three times with phosphate-buffered saline (PBS). For enhanced optical clarity and preservation prior to imaging, the fixed embryos were transferred to a 1:1 PBS:Glycerol solution and incubated overnight at 4°C. Subsequently, embryos were mounted on glass slides using glycerol-based mounting media (Glycerol + DABCO), covered with a 10 mm round glass coverslip, and the edges were sealed with transparent nail polish to prevent dehydration. Slides were stored at 4°C overnight to allow complete drying. Imaging was primarily performed using a confocal microscope (Zeiss, LSM 900).

Fluorescence intensity quantification was carried out using ImageJ software. For specific visualization of lipid droplets (LDs) within the larval liver, a 63x oil immersion objective was used, and fluorescence intensities were measured from random regions of interest (n=6) across the entire liver. To quantify LDs, the number of LDs per cell was counted from randomly selected cells (n=10) across the liver using ImageJ. These counts were then plotted for analysis.

To monitor the long-term distribution of lipids in the circulation, particularly within the tail vasculature of zebrafish larvae, Nile Red staining was performed. Six randomly selected larvae from each treatment group at various time points post incubation (6h, 12h, 24h, 48h, 120h) were fixed in 4% PFA overnight at 4°C and washed three times in PBS. Nile Red staining was conducted following (45), where larvae were immersed in a 0.79 mM Nile Red solution in embryo media and incubated in the dark at room temperature for 30 minutes. Excess dye was removed by three washes in PBS, and the stained larvae were then transferred to 50% glycerol in PBS and stored overnight at 4°C. Finally, larvae were mounted on transparent glass slides in 70% glycerol with DABCO for imaging. Confocal microscopy for Nile Red imaging was performed using the Zeiss confocal microscope (LSM 900) with excitation at 543 nm and emission at 598 nm. Consistent imaging parameters (laser intensity and exposure time) were maintained across all samples. Nile Red fluorescence intensity in the tail vasculature, indicative of circulating lipids, was quantified using ImageJ software and expressed as fold change relative to larvae treated with empty liposomes at each corresponding time point.

#### Effect of KTDP on Morphology, Mortality and Locomotion of Zebrafish Larvae

To evaluate the impact of KTDP on zebrafish larvae, individuals were subjected to different liposome treatments. For each experimental condition, zebrafish larvae (n=30) were placed in each well of a 6-well plate, with 3 ml of media added to each well. All experiments were performed in triplicate to ensure reproducibility. Larvae were incubated for 6 hours (at 30 RPM shaking, 28.5 ℃) with either empty egg liposomes (vehicle control), liposomes encapsulating KTDP, or liposomes encapsulating a KTDP-Mutant peptide, prepared as previously described. Following this 6-hour incubation period, larvae were transferred to fresh E3 medium and maintained under standard conditions with normal feeding protocols.

Morphological evaluations were conducted at 6, 12, 24, 48 and 120 hours post-incubation (hpi) with the liposome treatments. Prior to imaging, larvae were anesthetized in E3 medium containing 0.06% Tricaine. Brightfield images were captured using an stereomicroscope (Olympus MVX10). Consistent illumination, magnification, and exposure settings were maintained across all samples and time points. Qualitative assessments focused on gross morphological changes, including body curvature, edema formation (e.g., pericardial), developmental milestones (e.g., swim bladder inflation, yolk sac absorption), and overall structural integrity compared to the vehicle control group.

Survival rates were monitored daily for 120 hours post-incubation. Larvae were examined at 24-hour intervals, and mortality was recorded. Cumulative mortality percentages were calculated for each group at each time point. Survival curves were generated based on these data. Environmental conditions (28°C, 14:10 light:dark cycle) were strictly maintained throughout the assay period.

Escape response was assessed at 120 hpi as described (46) with some modifications. Individual larvae (n=10 per condition) were transferred to the center (Zone 0) of a 35 mm Petri dish containing E3 medium. The dish surface was conceptually divided into concentric zones (Diameter Zone 0: 7.78 mm, Zone 1: 16.85 mm, Zone 2: 25.92 mm, Zone 3: 35 mm). A standardized tactile stimulus was delivered to the caudal fin region using a fine micropipette tip to elicit an escape response. The zone in which the larva first ceased significant movement following the stimulus was recorded. The distribution of larvae across zones was plotted as the mean of their zone no. Spontaneous locomotor activity was assessed at 120 hours post-incubation (hpi). Briefly, individual larvae were placed in separate wells of a 6-well plate containing E3 medium. Larval movement was recorded for 3 minutes using a video camera (30 fps) mounted above an LED-illuminated surface supporting the plate. Video files were processed using ZebraZoom software (47) to automatically track larval position and quantify locomotion parameters. For comparative analysis, the mean total distance traveled (mm) and mean angular velocity (deg/s) for each treatment group were expressed as fold changes relative to the mean values obtained from the vehicle control group (empty liposomes). All behavioral recordings were conducted at 28°C under consistent lighting conditions to minimize external variability.

#### Animal Husbandry and Grouping for Adult Zebrafish Experiments

One-year-old adult zebrafish (Danio rerio) of both sexes were equally and randomly allocated into two experimental groups: (1) empty liposomes, and (2) KTDP-liposomes. Fish were housed in a custom-designed standalone recirculating aquaculture system. This system consisted of fish tanks placed on a draining tray connected to a lower reservoir with biological filters and a submersible pump. Water was continuously cycled back into the tanks via inlet tubing, and overflow was drained back to the reservoir, maintaining stable water quality and flow conditions. Water quality parameters such as temperature, pH, and ammonia levels were monitored regularly to ensure optimal husbandry conditions.

#### KTDP Administration via Oral Gavage in Adult Zebrafish

KTDP was encapsulated in egg-yolk liposomes and administered at a concentration of 250 µg/mL, identical to the dose used in larval studies. Each fish received 5 µL of liposomal solution once daily in the morning, followed by standard feeding for the rest of the day. Gavage was performed for three consecutive days.

Due to the technical challenges associated with oral gavaging in adult zebrafish, we adopted and modified the method described by Collymore et al. (2013). Briefly, a custom sponge block with a midline groove was used to immobilize the fish in a vertical orientation. A femtotip, trimmed to allow better flow, was fitted onto a 10 µL pipette and loaded with the liposomal dose. Fish were anesthetized in 100 µg/mL tricaine, placed in the moistened sponge with their head exposed, and the tip was gently inserted ∼3 cm into the oral cavity to dispense the solution. To prevent regurgitation, fish were kept vertically in the sponge for an additional 30 seconds post-gavage before being transferred to a recovery tank for 15 minutes. Any fish displaying bleeding, distress, or abnormal behavior were excluded from further experimentation.

#### Sample Collection from Adult Zebrafish

On the third day, after the final dosing, fish were fasted and sacrificed four hours post-gavage. Blood was collected from the tail vein as described by Gali et al. (2019) and Pedroso et al. (2012). Following anesthesia, fish were wiped with 70% ethanol, and the tail was severed just caudal to the anal fin using a sterile blade. Fish were then placed in a perforated 0.5 mL microcentrifuge tube, which was inserted into a 1.5 mL collection tube and centrifuged at 50 × g to collect blood. Serum was separated from the supernatant for further analysis. Liver and anterior gut tissues were dissected, washed in cold PBS, and imaged immediately under an Olympus stereomicroscope in FITC and TRITC channels. Tissues were then snap-frozen in liquid nitrogen and stored at −20°C for further biochemical analysis.

#### Molecular Dynamics Simulations

##### System setup

The initial PDB structure of the Kinesin Tail Domain protein (KTDP) is modeled in I-TASSER (48) due to the unavailability of experimentally reported structures in the Protein Data Bank. The modeled structure consisting of 110 residues, is solvated in explicit water and equilibrated for 1 ns in an isothermal-isobaric (NPT) ensemble. The final equilibrated protein structure is extracted for further modeling steps. Two membrane-protein systems are prepared to study the differential binding of KTDP with monolayer and bilayer cargo. A model monolayer system is constructed by extracting the PDB structures of 1,2-dioleoyl-sn-glycero-3-phosphocholine (DOPC), 1,2-dioleoyl-sn-glycero-3-phosphate (DOPA), and Triacylglycerols (TG) molecules from membrane patches built in CHARMM-GUI (49, 50). PACKMOL (51) is used to build the model monolayer system by defining three boxes at different square planes with of sides measuring ca. 12.3 nm. The planar surfaces are designated to the membrane leaflets consisting of DOPC and DOPA (first and third) in a 95:5 ratio and a neutral lipid layer of TGs (second). The upper and lower leaflet consisting of DOPC and DOPA is added to support the middle TG layer without compromising the atomic resolution and stability. PACKMOL-generated monolayer PDB structure is solvated with the TIP3P water model and equilibrated for 400 ns at 303 K in the isothermal-isobaric (NPT) ensemble. The final PDB structure is taken as the initial monolayer configuration interacting with the KTDP. The bilayer system (devoid of TG) is generated within the CHARMM-GUI server. The center of mass (COM) of KTDP is kept ca. 2.5 nm apart from the COM of phosphate of upper leaflet in both systems to ensure unbiased interaction. Two 6.4 nm thick water layers are introduced at both sides of the leaflets in the monolayer, whereas in the bilayer, the water thickness is 5 nm at each side. The final monolayer and bilayer consisted of 299888 and 196670 atoms, respectively. The number of molecules in monolayer and bilayer systems are :-

Monolayer DOPC - 418 (209 in each leaflet), DOPA - 22 (11 in each leaflet), TG - 300, Water - 62577, Ions - 9 (Na+ to neutralize the system).

Bilayer DOPC - 418 (209 in each leaflet), DOPA - 22 (11 in each leaflet), Water - 44871, Ions - 9 (Na+ to neutralize the system).

Fully atomistic molecular dynamics (MD) simulations are performed with the GROMACS 21.1 MD engine (52) with the CHARMM36m forcefield (53). TIP3P explicit water model is used for solvation. Bond lengths involving hydrogen atoms are held fixed using the LINCS algorithm (54). A non-bonded cutoff of 1.2 nm is used, and long-range electrostatic interactions are computed with the Particle Mesh Ewald (PME) method (55). The systems are initially minimized with the steepest descent algorithm, The system is equilibrated for 5 ns at a constant temperature of 310 K using the Nose-Hoover thermostat and an isobaric and semi-isotopic pressure of 1 bar using the Parrinello-Rahman barostat. Velocities are periodically reassigned over 1 μs, followed by production runs of 0.25 μs. Effective trajectories of 1.25 μs were generated for each system with a timestep of 2.0 fs. ChimeraX is used for visualization (56).

##### Analysis

Area per lipid (APL) is defined as the accessible leaflet area per lipid molecule. A higher APL value represents a higher accessible area for a lipid, corresponding to less lipid packing (57, 58). the standardized FATSLiM algorithm (59) was used for the APL.

Lipid occupancy (O*ij*, where *i* = lipid type ; *j* = peptide residue) is defined by the time fraction in which a particular amino acid residue forms contact with a lipid molecule, and is estimated with a cutoff distance of 0.7 nm. This calculation is done by using the PyLipID toolkit [20]. For the *i*^th^ lipid (DOPC or DOPA) and *j*^th^ peptide esidue, the ΔO*ij* is calculated as the following,

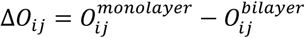

The average DOPC-DOPA occupancy of a KTDP residue (= <O>*j*) is determined by averaging the DOPC and DOPA lipid occupancies of each KTDP residue for monolayer and bilayer systems.

Lipid fraction per residue (*Lf*) is determined as 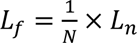. Here *N* is the number of residues interacting with *Ln* lipids within a cutoff of 0.7 nm. Interactions with the headgroups (for DOPC and DOPA) or with TG molecules were considered. of each residue having favorable interaction in monolayer and bilayer. The smooth probability distribution envelope *P(Lf)* was obtained from the discrete data with Kernel Density Estimation (KDE).

## Supporting information

MD Simulation KTDP-Bilayer

MD Simulation KTDP-Monolayer

## ACKNOWLEDGEMENTS

RM acknowledges funding from IIT Bombay and through Senior Fellowships from the Department of Biotechnology – Wellcome Trust India Alliance (Grants IA/S/11/2500255 and IA/S/19/2/504634). SSK acknowledges a SwarnaJayanti Fellowship from SERB, Govt. of India (SB/SJF/2021–22/01). We thank the Tata Institute of Fundamental Research for animals and access to their animal facility. H.C. acknowledges the Prime Minister’s Research Fellowship (PMRF) for Ph.D. Authors acknowledge the National Supercomputing Mission (NSM) for providing computing resources of “PARAM Shakti” at IIT Kharagpur, and “PARAM Rudra” at S. N. Bose National Center for Basic Sciences (SNBNCBS) which are implemented by C-DAC and supported by the Ministry of Electronics and Information Technology (MeitY) and Department of Science and Technology (DST), Government of India. N.S acknowledges the STARS-2 Grant (2023-0210) from Ministry of Education received as co-PI (with Arnab Gupta), and the ANRF (formerly SERB; Grant No. CRG/2020/005610), for computational equipment.

## Abbreviations

(APL): Area per lipid
(ALDs): Artificial Lipid Droplets
(CE): Cholesterol Esters
(ER): Endoplasmic Reticulum
(GFP): Green Fluorescent Protein
(KHC): Kinesin Heavy Chains
(KLC): Kinesin Light Chain
(KTD): Kinesin Tail Domain
(KTDP): Kinesin Tail Domain Peptide
(LD): Lipid Droplet
(MD): Molecular Dynamics
(PA): Phosphatidic Acid
(PC): Phosphatidylcholine
(RMSD): Root mean square deviation
(SS): Secondary structure
(TG): Triacylglycerol
(VLDL): Very Low Density Lipoproteins

## SUPPLEMENTARY FIGURES AND SUPPLEMENTARY MOVIE CAPTIONS TRIPATHY ET AL

**SUPPLEMENTARY FIGURE S1.**
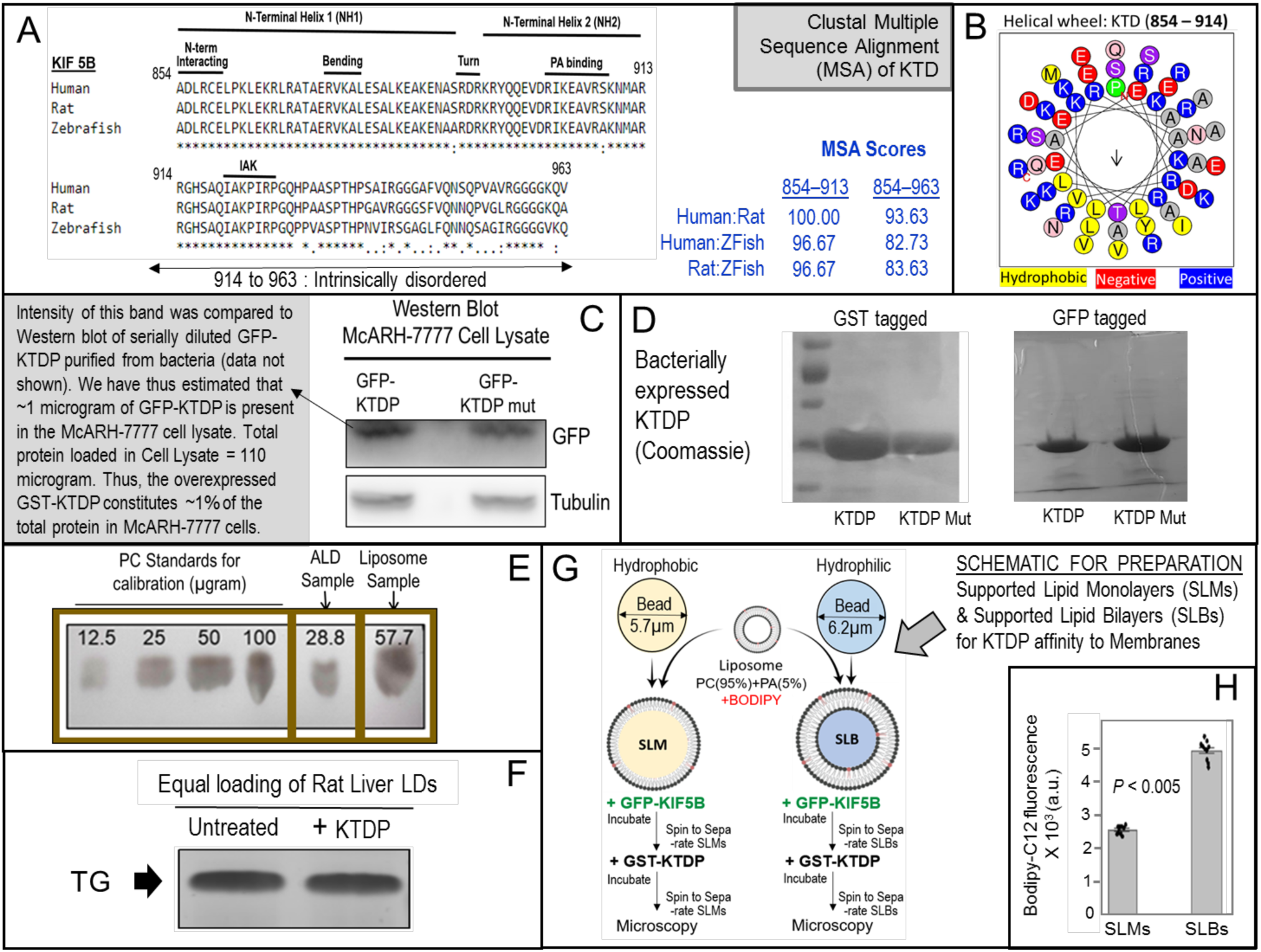
Details related to KTDP, Controls for *In-vitro* experiments and Schematic of SLM and SLB Experiments. A. Clustal multiple sequence alignment of KTD from KIF5B (Kinesin-1) across Human, Rat and Zebrafish species. The MSA scores are mentioned. High conservation is seen in the 854-913 AA region that has coiled coil propensity and includes the PA-binding region (901-909). B. Helical wheel alignment of KTD AA 854-914 shows separation of hydrophobic and charged residues to opposite faces of the helix, suggesting an ability to form amphipathic helices upon membrane interaction. C. Western blot of GFP-KTDP and GFP-KTDP-Mutant after overexpression in McARH-7777 cells. D. Expression of GFP and GST-tagged KTDP and KTDP-Mutant in bacterial systems. E. Liposomes and ALDs were prepared using (PC 95% + PA 5%). A thin layer chromatograph (TLC) was run using known dilutions of PC along with the liposome and ALD samples. The estimated amount of PC in liposome sample and ALD sample (as obtained using the PC standards) is mentioned. These values of PC were used to normalize the liposome and ALD samples, considering that 95% of both ALD and liposome membranes consists of PC (see main text). F. Thin layer chromatograph (TLC) to detect TG shows equal loading of LDs (purified from rat liver) that were left untreated, or were treated with KTDP. G. Schematic to explain the supported lipid monolayer (SLM) and supported lipid bilayer (SLB) assays for determining affinity of KTDP to monolayer and bilayer membranes. H. Twice the amount of BODIPY fluorescence is detected on SLBs (bilayer) as compared to SLMs (monolayer). Each data point represents the integrated fluorescence measured along a circular profile around individual SLMs or SLBs. Errors are SEM.

**SUPPLEMENTARY FIGURE S2.**
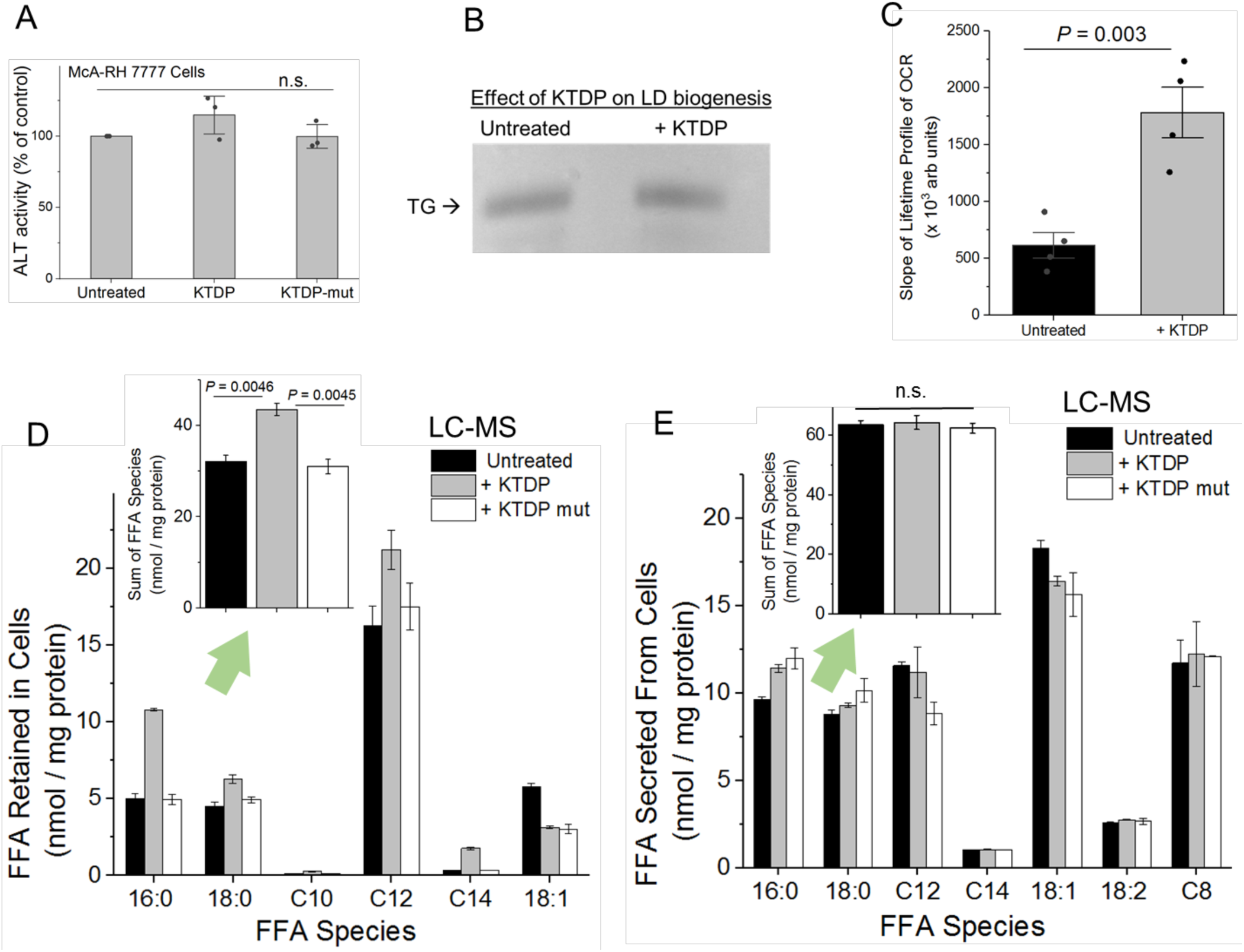
ALT activity, LD biogenesis, OCR and FFA measurements using Mc-ARH 7777 Cells. A. ALT activity in McA-RH 7777 cells that are untreated, overexpressing KTDP or KTDP-mutant. Data are mean ± SEM; ns = not significant B. TLC of McA-RH 7777 cells that are untreated or overexpressing KTDP. Cells were first depleted of LDs by keeping in serum-free medium followed by addition of Oleic acid to induce LD biogenesis for 12 hours. Cells were then lysed, and TLC of cell lysate was run. The TG band is similar across conditions suggesting no effect of KTDP on LD biogenesis. C. Cellular oxygen consumption rate (OCR) in McA-RH 7777 cells showing increased cellular respiration in KTDP-overexpressing cells compared to untreated cells. Data are mean ± SEM. D. LC-MS measurement of free fatty acids (FFAs) retained in Mc-ARH 7777 cells. The sum of detected FFA species is also shown (errors have been propagated). There is a significant increase of FFAs inside cells after overexpression of KTDP, suggesting the activation of a lipolytic pathway. Errors are SEM. E. LC-MS measurement of FFA species secreted from Mc-ARH 7777 cells. The sum of detected FFA species is also shown (errors have been propagated). There is no significant effect of overexpressing KTDP on FFA secretion from cells. Errors are SEM.

**SUPPLEMENTARY FIGURE S3.**
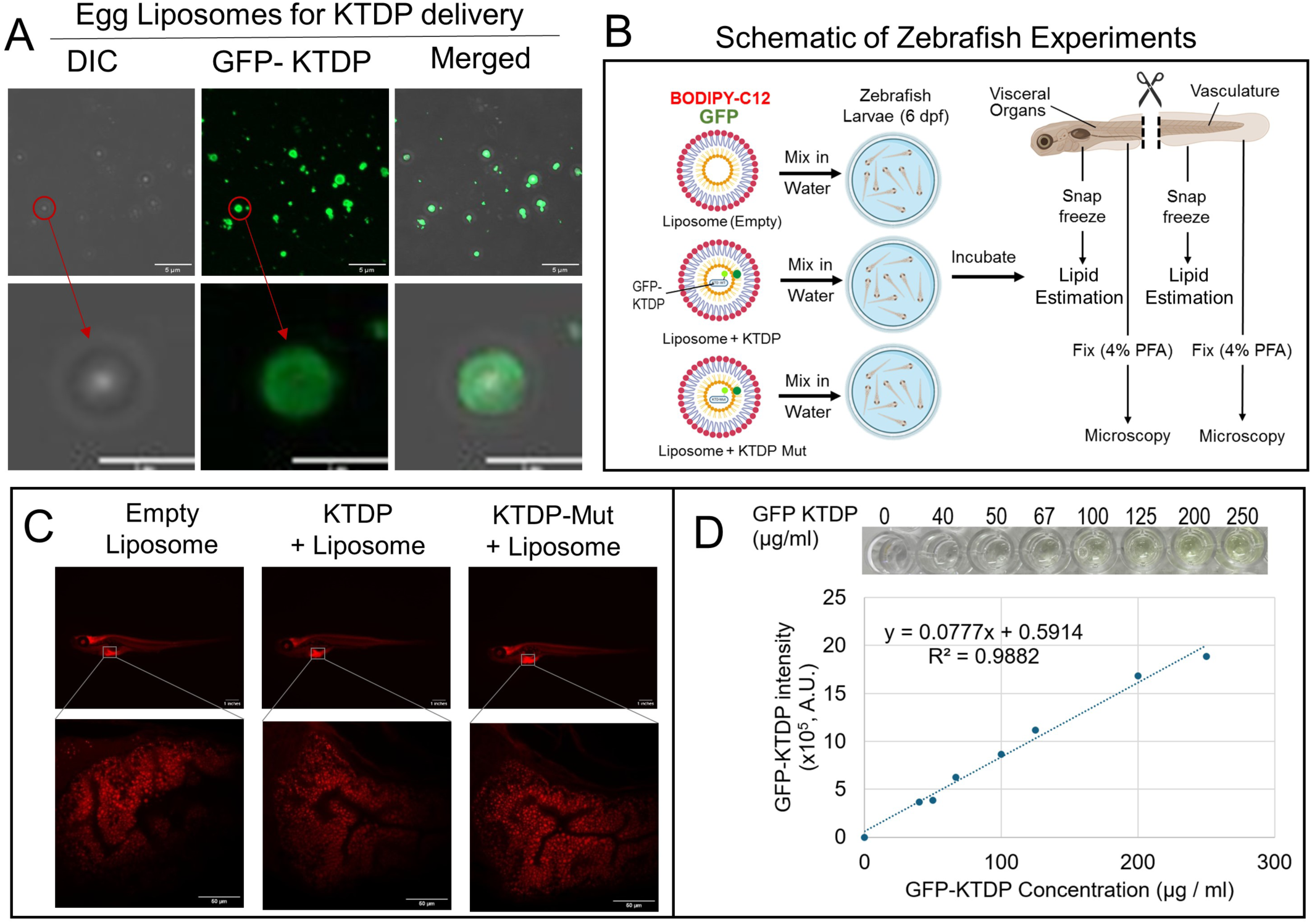
Experiments related to Egg-Liposomal Delivery KTDP in Zebrafish Larvae. A. Microscopy of Egg liposomes containing GFP-KTD. DIC (Differential Interference Contrast) and Confocal images are shown. GFP signal is clearly visible within nearly all liposomal vesicles. Scale bar = 5 μm. B. Schematic representation of the feeding protocol in which BODIPY-C12-liposomes (with and without KTDP) are administered to 6 days post-fertilization (dpf) zebrafish larvae, followed by microscopy and biochemical analysis to observe the effect of KTDP on lipid secretion. C. Confocal images showing BODIPY fluorescence (red) in zebrafish larvae (6 dpf) and in the gut of larvae after incubation with BODIPY-containing liposomes across conditions (mentioned). Results indicate equal feeding across experimental groups. D. Fluorescence intensity gradient of GFP-KTDP corresponding to increasing protein concentrations. Standard curve showing the linear relationship between fluorescence intensity and protein concentration of purified GFP-KTDP.

**SUPPLEMENTARY FIGURE S4.**
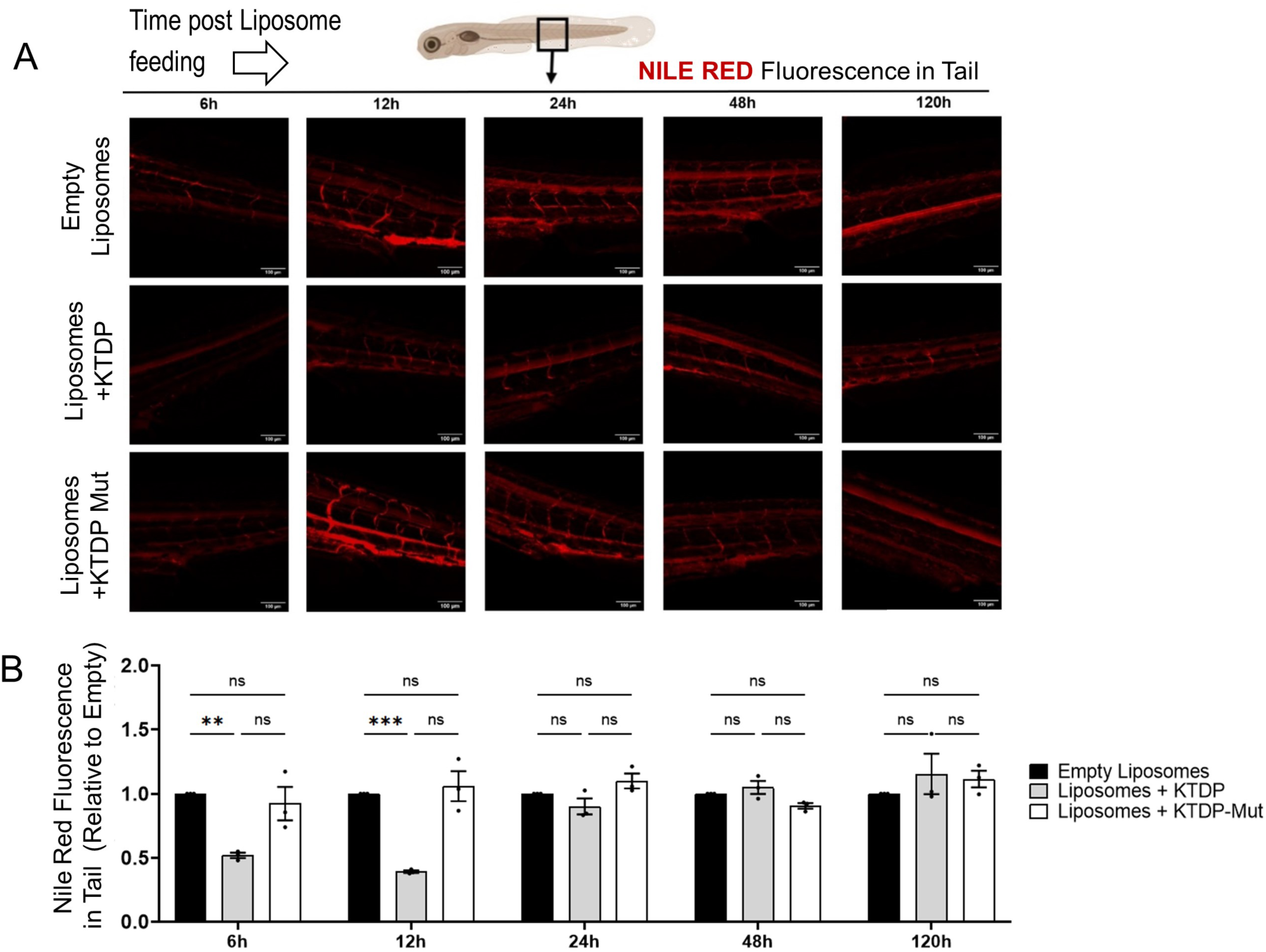
Time course of Lipids in Tail of Zebrafish Larvae after Liposomal feeding of KTDP. A. Representative confocal images of Tail region (vasculature) of Nile red stained Zebrafish larvae at 6, 12, 24, 48 and 120 hours post incubation with Egg Liposomes (unlabelled) that were Empty, containing KTDP or KTDP-mutant. B. Quantitative analysis of fluorescent intensity, expressed as fold change relative to larvae treated with Empty liposomes at that time-point. The data reveal ∼50% reduction of lipid content in the tail at 6 and 12 hours, followed by reversal to baseline levels. Overall fluorescence intensity at the 6 hour time point is lower because the lipids contained in egg-liposomes have presumably not yet been metabolized and secreted out into vasculature. *** denotes *P* ≤ 0.001, ** denotes *P* ≤ 0.01 and n.s. implies *P* > 0.05.

**SUPPLEMENTARY FIGURE S5.**
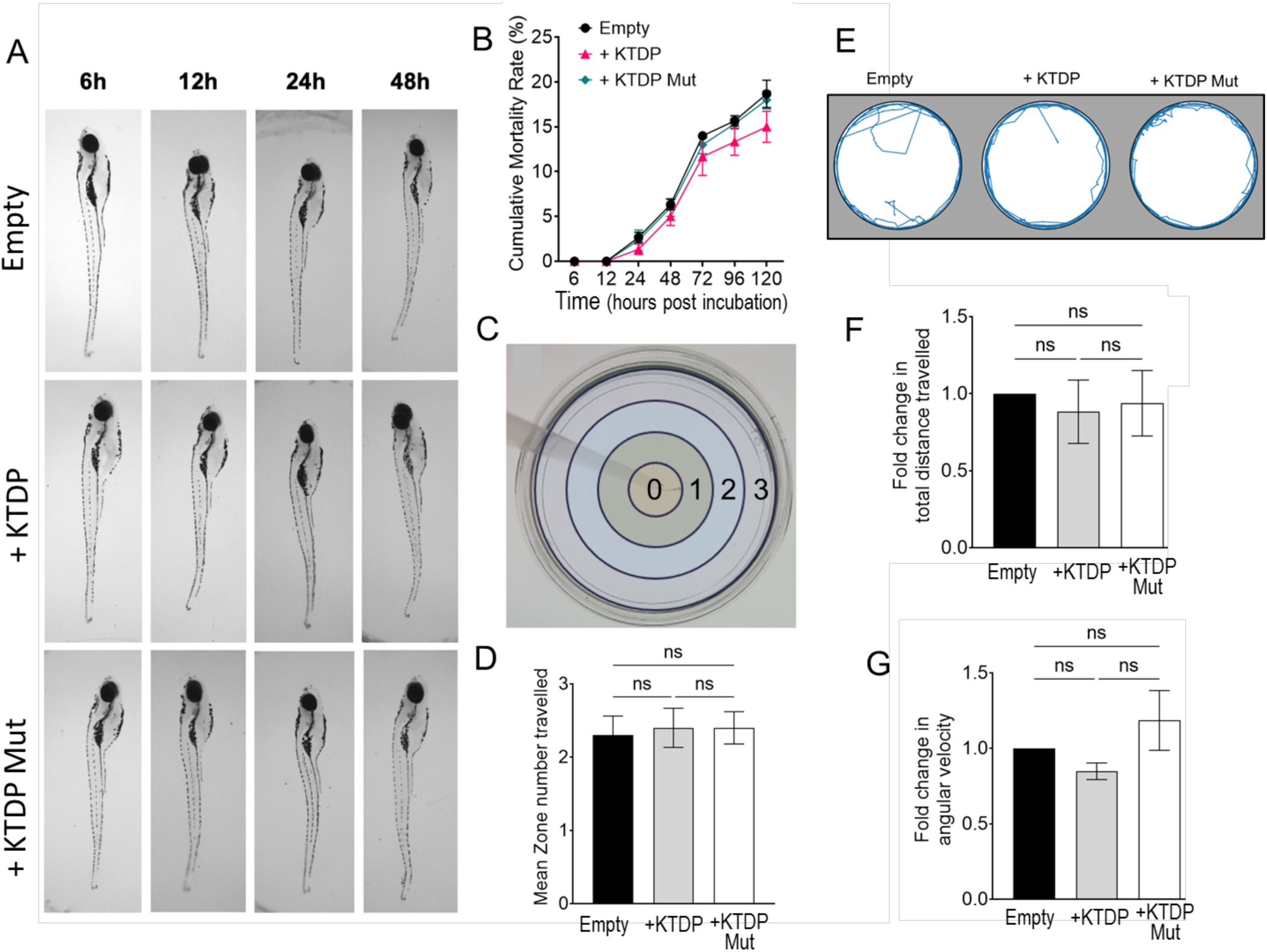
Effect of KTDP on Morphology, Mortality and Locomotion of Zebrafish Larvae. A. Images of Zebrafish larvae at 6, 12, 24, and 48 hours post-incubation with egg-liposomes that were Empty, containing KTDP or KTDP-Mut. No observable effect of KTDP is seen on the morphology of larvae. B. Mortality rates of larvae were monitored up to 120 hours post-incubation with egg-liposomes that were Empty, containing KTDP or KTDP-Mut. No significant differences are seen across experimental groups. C. Petri dish setup illustrating concentric zones for studying larval locomotion. Individual larvae were placed at the center and stimulated with a micropipette tip. The zone where individual larvae first stopped after stimulation was noted. Zones were defined concentrically, with the center being Zone 0 and the outermost being Zone 3. D. Results of micropipette stimulation indicate no significant differences in the first stop zones among experimental groups at 120 hours post KTDP feeding, suggesting no impairment in touch-evoked response (TER). E. Trajectories of individual larvae recorded over six minutes in a six-well plate that were analyzed using ZebraZoom software. F. Fold change in total distance traveled by larvae from video recordings. G. Fold change in angular velocity derived from the same video recordings. These results indicate no significant change in locomotory behavior after incubation with KTDP.

**SUPPLEMENTARY FIGURE S6.**
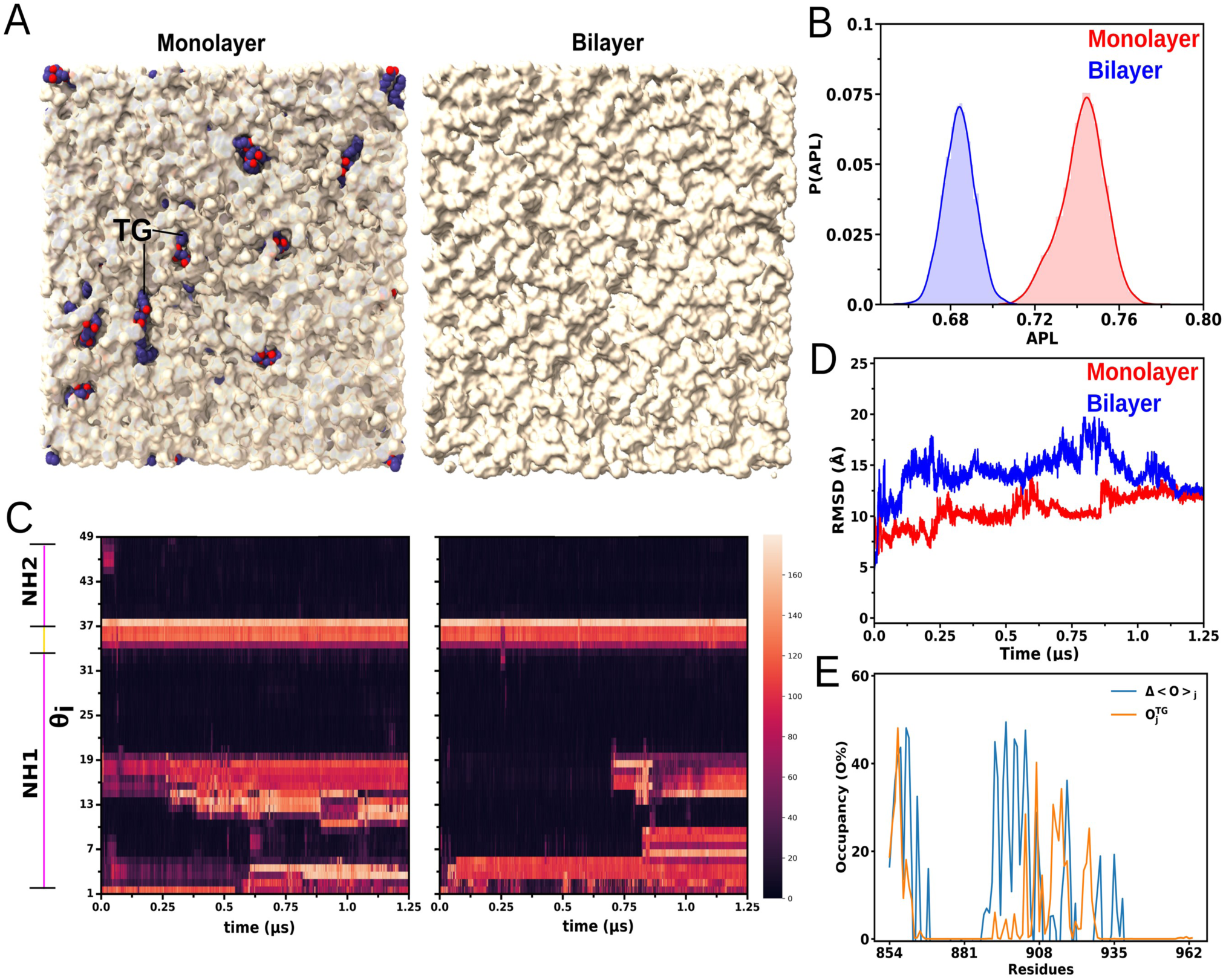
Additional Details for MD Simulations of KTDP interacting with Monolayer and Bilayer Membranes. A. A top view representation of membrane leaflets of monolayer and bilayer systems. Surface-exposed TG molecules (dark purple) are observed only on the monolayer. B. Probability distributions of the area per lipid (APL, in nm^2^) on monolayer and bilayer systems. C. Helix distortion of KTDP on the monolayer and bilayer systems as a function of time (see main text). The KTDP N-terminal helices (NH1 and NH2) are indicated. D. Backbone RMSD of KTDP in monolayer and bilayer over the simulation trajectory. E. Monolayer favouring average lipid occupancy (Δ< *O* >*_ij_*) > 0 and TG occupancy 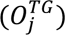 during KTDP-monolayer interactions. Note the correlation between TG and PC/PA interactions, suggesting that exposed TG patches on the monolayer (see panel A) are important for these interactions.

## SUPPLEMENTARY MOVIE CAPTIONS, TRIPATHY ET AL

### SUPPLEMENTARY MOVIE 1

Video showing the interaction of Kinesin tail domain peptide (KTDP) with Bilayer Membrane over a period of 125 microseconds. This video is based on MD simulations (see main text).

### SUPPLEMENTARY MOVIE 2

Video showing the interaction of Kinesin tail domain peptide (KTDP) with Monolayer Membrane over a period of 125 microseconds. This video is based on MD simulations (see main text).

